# De novo missense variants in TRIM28 identified in individuals with neurodevelopmental delay show features of transposable element activation

**DOI:** 10.64898/2026.07.02.735854

**Authors:** Laura Castilla-Vallmanya, Ninoslav Pandiloski, Carrie Davis-Hansson, Fereshteh Dorahezi, Ofelia Karlsson, Maria Isabel Alvarez-Mora, Irene Madrigal, Francisco Barros-Angueira, Alison M. Muir, Chitra Prasad, Samantha Colaiacovo, Christopher Douse, Susanna Balcells, Raquel Rabionet, Johan Jakobsson

**Affiliations:** Laboratory of Molecular Neurogenetics, Department of Experimental Medical Science, Wallenberg Neuroscience Center and Lund Stem Cell Center, BMC A11, Lund University; 221 84 Lund, Sweden; Laboratory of Epigenetics and Chromatin Dynamics, Department of Experimental Medical Science and Lund Stem Cell Center, BMC A11, Lund University; 221 84 Lund, Sweden; Biochemistry and Molecular Genetics Department, Hospital Clinic of Barcelona, IDIBAPS (Institut de Investigacions Biomèdiques August Pi I Sunyer); 08036 Barcelona, Spain; CIBER of Rare Diseases (CIBERER), Instituto Salud Carlos III; 28029 Madrid, Spain; Fundación Pública Galega de Medicina Xenómica; Santiago de Compostela, A Coruña, Spain; GeneDx LLC; Gaithersburg, MD, USA; Division of Genetics, Department of Pediatrics, London Health Sciences Centre, Western University London; Ontario, Canada; Department of Genetics, Microbiology and Statistics, Facultat de Biologia, Universitat de Barcelona; 08028 Barcelona, Spain; Institute of Biomedicine of the Universitat de Barcelona (IBUB); 08028 Barcelona, Spain; Institut de Recerca Sant Joan de Déu (IRSJD); Barcelona, Spain

## Abstract

TRIM28 is an epigenetic co-repressor protein that silences transposable elements (TEs). Although loss-of-function studies in mice led to neurodevelopmental defects, a functional link between TRIM28 and human neurodevelopment has yet to be established. In this study, we describe two patients with neurodevelopmental delay who carry *de novo* TRIM28 missense variants. Using CRISPR-edited induced pluripotent stem cell lines and differentiated neural organoids, we demonstrate that these variants result in the loss of the histone mark H3K9me3 over TEs. This releases the regulatory potential of TEs resulting in altered expression of nearby genes. These findings could be replicated using CRISPRi-based TRIM28 silencing, which suggests that the two variants result in a loss of function. Our results highlight the critical role of TRIM28 in regulating TEs during human brain development, establishing a link between *TRIM28* variants and neurodevelopmental delay.

**One Sentence Summary:** TRIM28 variants disrupt epigenetic control of transposable elements in the developing human brain, linking them to neurodevelopmental delay.

## INTRODUCTION

Transposable elements (TEs) are mobile, repetitive sequences that comprise 50% or more of the human genome^1–4^. The majority of human TEs are retrotransposons, which are remnants of ancient transposition events that have become fixed in the human genome. Only a small percentage of retrotransposons remain capable of transposition^5–7^. However, many TEs carry sequences that have the potential to influence gene expression^8,9^. It is becoming increasingly clear that TEs can act as gene regulatory elements that control and fine-tune gene expression. For instance, we and others have demonstrated that TEs play an essential role in gene regulatory networks in human neural cells by acting as alternative promoters, enhancers, repressors, and non-coding regulatory RNAs^10–15^. Consistent with their important role in gene regulatory networks, aberrant TE activation has been associated with various human diseases^16–18^. Nevertheless, the role of TEs in human physiology and disease remains largely unexplored.

The epigenetic co-repressor protein Tripartite Motif Containing 28 (TRIM28), also known as KAP1 or TIF1β, is a master regulator of TEs^19,20^. TRIM28 is recruited to target TE sequences through its association with Krüppel-associated box domain zinc finger proteins (KZFPs)^21–24^. There, TRIM28 serves as a scaffold protein for a complex of repressive proteins including the histone methyltransferase SETDB1, the nucleosome remodelling and deacetylation (NuRD) complex, heterochromatin protein 1 (HP1) and DNA methyltransferases (DNMTs)^25–27^. The TRIM28 protein is an antiparallel homodimer with an asymmetry at the C-terminal domain and multiple SUMOylation and phosphorylation sites^28–30^. These properties functionally impact TRIM28 interactions as only one of the two putative binding sites may be occupied by the KRAB-ZFPs, HP1 and probably also the rest of its interactors. This repressive complex orchestrates the establishment of local heterochromatin at TEs, characterised by the presence of H3K9me3, that limit TE expression and their regulatory impact on the genome. In line with this, genetic ablation of *TRIM28* in mammalian cells leads to transcriptional activation of TEs and reveals their potential as gene regulatory elements with a capacity to alter the expression of nearby genes^31–33^.

Evidence from mouse models shows that TRIM28 plays a critical role in brain development. We have previously shown that deletion of *Trim28* during postnatal forebrain development or heterozygous deletion of *Trim28* during early brain development results in complex behavioural changes, including hyperactivity and impaired stress response^34,35^. Additionally, heterozygous germline deletion of *Trim28* in mice has been shown to induce abnormal exploratory behaviour^36^. These findings demonstrate that disruption of Trim28 levels in the mouse brain results in behavioural changes, similar to those observed in neurodevelopmental and psychiatric disorders^37,38^. However, the relevance of these observations to humans remains unclear, as *TRIM28* variants or mutations have not yet been associated with neurodevelopmental phenotypes.

In this study, we describe two individuals with neurodevelopmental delay that carry *de novo* heterozygous missense variants in *TRIM28*. Using gene-edited human induced pluripotent stem cell (hiPSC)-derived neural organoids, we found that cells expressing these TRIM28 variants exhibited a loss of H3K9me3-covered heterochromatin at TEs. This phenomenon could be replicated by TRIM28 loss-of-function experiments. The loss of heterochromatin at TEs unleashed their regulatory potential, disrupting gene regulatory networks and resulting in the aberrant expression of genes located near TRIM28-controlled TEs. Many of these TRIM28/TE-controlled genes are relevant to early brain development and neurodevelopmental disorders. In summary, our results suggest the critical importance of controlling TE activity in human brain development and link *de novo* TRIM28 missense variants to neurodevelopmental delay.

## RESULTS

### De novo TRIM28 variant carriers display a neurodevelopmental phenotype

Using exome sequencing (ES) we found heterozygous variants in *TRIM28* in two unrelated boys presenting with neurodevelopmental impairments. Both variants arose *de novo* and consisted of missense substitutions at codons Pro654Leu (c.1961C>T; p.P654L) and Leu708Pro (c.2123T>C; p.L708P) (Fig 1a). Segregation was confirmed by familial exome and/or Sanger sequencing (Fig. S1). Both variants map to the C-terminal PHD (plant homeo-domain) tandem bromodomain (PHD-Bromo) region of the TRIM28 protein (Fig. 1b). Clinically, both individuals presented with moderate intellectual disability, and autism spectrum disorder, with diagnosis established at 31 months and 10 years old. Phenotypic findings are summarized in Table 1.

**Fig. 1.**
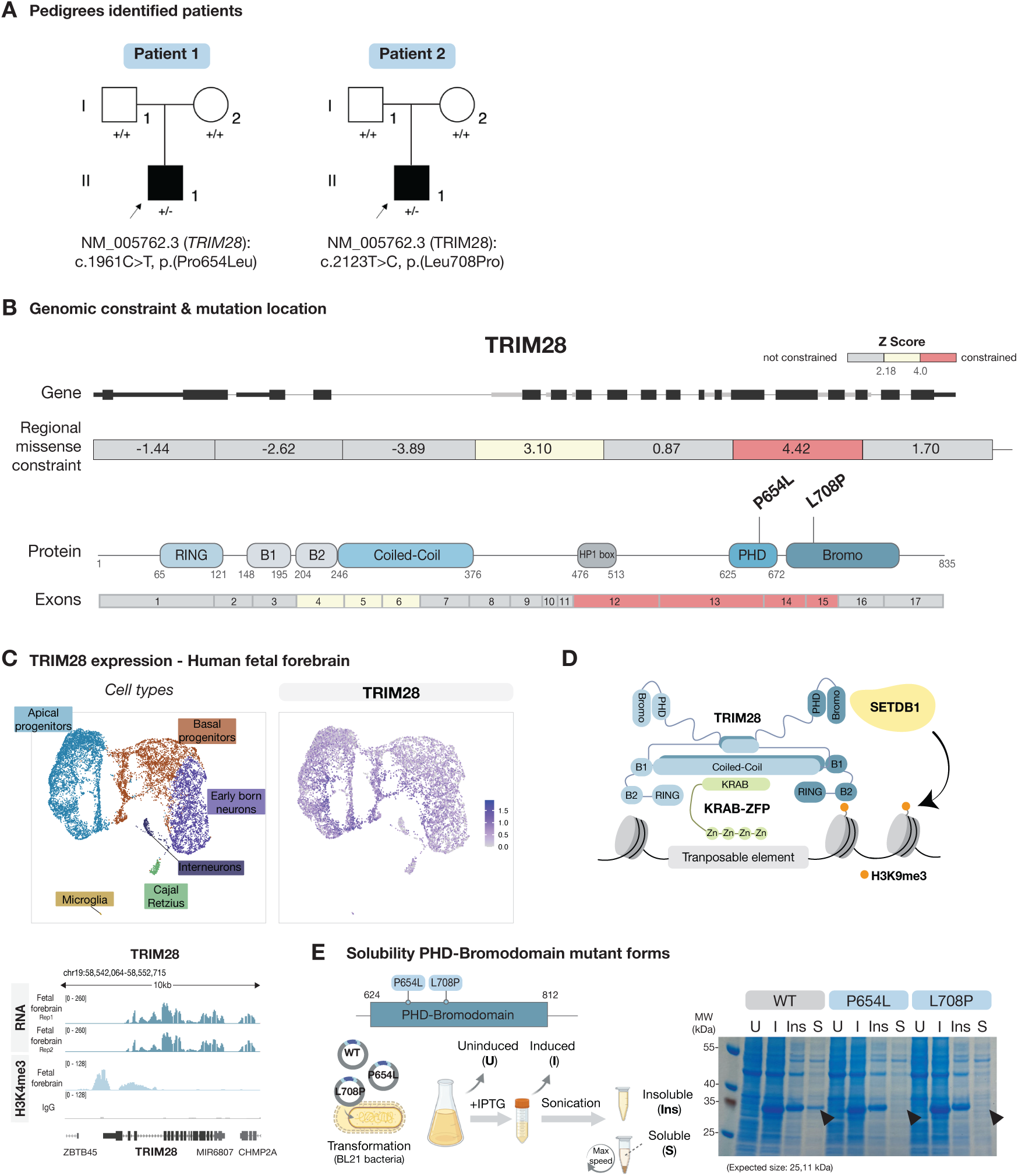
The identified *TRIM28* variants in the patients have pathogenic potential. (**A**) Pedigrees showing the *de novo* variants in *TRIM28* identified in patients via whole exome sequencing. (**B**) Top: Gene schematics and regional 1 kb missense constraint scores from GnomAD^39^. Bottom: Schematics of the functional protein domains of TRIM28 and corresponding exons coloured by regional missense constraint score. The locations of the identified variants in the patients are indicated. (**C**) Top: UMAP of snRNA-seq data showing *TRIM28* expression in 7.5-10.5 weeks human fetal forebrain tissue. Bottom: Genome browser tracks showing the bulk RNA expression of *TRIM28* with an H3K4me3 peak on its promoter in 10-10.5 weeks human fetal forebrain tissue. Data from Garza et al., 2023^40^. (**D**) Model of TRIM28-mediated H3K9me3 deposition over TEs. Based on Stoll et al., 2019^28^ and Stoll et al., 2022^30^. (**E**) Left: Schematics of construct and experimental design followed to evaluate solubility of wild-type (WT) and mutant forms of the PHD-bromodomain. Right: Coomassie-stained protein gel; U: uninduced; I: Induced; Ins: Insoluble Fraction; S: Soluble Fraction.

**Table 1.** Detailed genetic and phenotypic characterization of patients included in this study.

|  | Patient 1 | Patient 2 |
| --- | --- | --- |
| Age of diagnosis | 2y9m | 10y |
| Age at last evaluation | 17 y | 21y |
| Sex | Male | Male |
| **Variants** |  |  |
| Genomic variant (GRCh38) | chr19:58549629 C>T | chr19:58549965 T>C |
| Inheritance mode | de novo | de novo |
| cDNA (NM_005762) | c.1961C>T | c.2123T>C |
| Protein | p.(Pro654Leu); | p.(Leu708Pro); |
| Localization in TRIM28 | PHD Domain | Bromodomain |
| Regional constraint (gnomad 1kb windows) | Zscore 4.42 \| o/e 0.74 | Zscore 4.42 \| o/e 0.74 |
| Alphamissense prediction | 0.9911 | 0.9993 |
| REVEL | 0.795 | 0.483 |
| CADD score (V1.7) | 32 | 25.7 |
| **Biometry at birth** |  |  |
| Gestational age | 36w | 39 w |
| Weight (Kg) (SD) | 2.690 Kg (p8, -1.44DE) | 4.3Kg |
| Length (cm) (SD) | 51 cm (p70, 0.54 DE) | NA |
| Postnatal concerns | Hemorrhagic disease of newborn; Jaundice | Pyloric stenosis |
| **Biometry at diagnosis** |  |  |
| Age | 31 months | 10y |
| Weight | 14Kg (50 perc) | 29Kg (10perc) |
| Height | 100cm (>95 perc.) | 132 cm (<3 perc).<br>Treated with growth hormone |
| Head circumference | 51cm (85 perc) | 54.5cm (50 perc) |
| **Biometry at last evaluation** |  |  |
| Age | 17y |  |
| Weight | 86.6 Kg (p93, 1.52 DE) |  |
| Height | 169.8 cm (P22, -0.78 DE) |  |
| **Development** |  |  |
| Motor delay | Yes |  |
| started walking at | 18 months | 13 months |
| Poor gross motor coordination | Yes | Yes |
| Psychomotor retardation | Yes | Yes |
| Speech delay | Yes | Yes |
| started talking at | 2-3 years (poor comprehension) | 3-4 years (poor comprehension) |
| Global developmental delay | Yes | Yes |
| Intellectual disability | Moderate ID | Moderate ID |
| Autism Spectrum Traits | Yes | Yes |
| Behaviour Phenotype (other) | NA | Significant problems, violent behaviour, anxious behaviour; Tic disorder |
| **Neurologic features** |  |  |
| Hypotonia | Yes | -- |
| Seizures | No | Febrile seizures |
| Structural findings on neuroimaging | RMC without alterations | Nonspecific white matter changes without progression |
| **Dysmorphic features** |  |  |
| Microcephaly | No | No |
| Other skull shape anomalies | No | No |
| Ptosis | No | No |
| Blepharophimosis | No | No |
| Epicanthal folds | No | Mild |
| Hypertelorism | No | Yes |
| Abnormally set or dysplastic ears | No | Slightly low set ears |
| Abnormality of the nose | - | Broad nasal bridge |
| Abnormality of the palate/mouth | - | Carp like mouth, prominent lower lip |
| Abnormality of the nipples | - | Right nipple flat |
| Cryptorchidism | - | Undescended left testicle |
| **Other findings** |  |  |
| Fetal finger pads | - | Yes |
| Psoriasiform dermatitis | Yes | -- |
| Abnormalities of the digestive system | - | Constipation |
| Kidneys | - | Nephrocalcinosis |
| **Oncological findings** |  |  |
| Wilms tumor | No | No |

According to bulk gene expression data from GTEx (GTEx Analysis Release V10, dbGaP Accession phs000424.v10.p2)^39^, *TRIM28* is ubiquitously expressed in adult human tissues. Using our previously published bulk-RNAseq, snRNA-seq and CUT&RUN data from fetal forebrain samples aged 7.5 to 10.5 weeks after conception^40^, we confirmed that *TRIM28* is highly expressed in the developing human forebrain (Fig. 1c). H3K4me3 CUT&RUN analysis confirmed that the *TRIM28* promoter is active in this tissue (Fig. 1c).

According to GnomAD constraint data, *TRIM28* is slightly intolerant to missense mutations (Z = 1.56) and highly intolerant to loss-of-function (LOF) variants (probability of being loss-of-function intolerant = 1, loss-of-function observed/expected fraction = 0.09) (gnomAD v.4.1.0, https://gnomad.broadinstitute.org/)^41^. Notably, the PHD-bromodomain, located at the 3’ end of TRIM28 and where the p.Pro654Leu and p.Leu708Pro variants are present, is highly intolerant to variation (Z=4.42) (Fig. 1b). The two putative pathological variants are absent in the general population (gnomAD v.4.1.0)^41^, consistent with a potential functional impact. According to MetaRNN^42^ the p.Pro654Leu and p.Leu708Pro variants are classified as strongly pathogenic. The PHD domain directs SUMO conjugation of an adjacent bromodomain, acting as an intramolecular SUMO E3 ligase^43^. When recruited to a specific genomic location by a KRAB-ZFP, the SUMO-modified TRIM28 is able to bind SETDB1, which is required for H3K9me3 deposition, heterochromatin formation and transcriptional silencing^25,27,28,43,44^ (Fig 1d).

To investigate whether the p.Pro654Leu and p.Leu708Pro variants are structurally disruptive and affect the proper folding of the PHD-bromodomain, we produced the wild-type (WT) and mutant forms of this protein domain (p.P654L and p.L708P) in a bacterial overexpression system. We found that the WT protein was present in the soluble fraction, while the mutant forms were completely insoluble, suggesting that both p.P654L and p.L708P variants result in misfolding of the TRIM28 PHD-bromodomain (Fig.1e).

### Generation of TRIM28 mutant and loss-of-function iPSCs and neural organoids

To investigate if the p.P654L and p.L708P missense variants cause loss of TRIM28 protein function we used CRISPR/Cas9 gene editing to generate hiPSCs harbouring the TRIM28 mutations at homozygosity (n=2 lines for each mutation, Fig. 2a&b). We used hiPSCs that had undergone the same clonal selection procedure as the CRISPR mutants as control cell lines for the P654L and L708P mutant lines (Fig 2a). In parallel, we generated hiPSCs with silenced TRIM28 expression using lentiviral-based CRISPR interference (TRIM28-CRISPRi, 2 independent gRNAs, Fig 2a&c). For the CRISPRi control, we used a gRNA that targeted a sequence not present in the human genome (Fig 2a). The CRISPR-edited hiPSC lines displayed normal colony morphology and appropriate levels of pluripotency marker genes, such as Nanog and OCT4, with no evidence for expression of genes linked to differentiation (Fig 2d & Fig. S2a, b). The edited clones did not show chromosomal abnormalities (Fig. S2c) nor signs of on-target hemizygosity (Fig. S2d). The missense mutations did not affect *TRIM28* mRNA levels (Sup S2e) or the nuclear subcellular localization of the TRIM28 protein (Fig. S2f), though TRIM28 protein levels were reduced in both mutant iPSCs, particularly in the P654L clones (Fig. S2g). This may be due to increased protein degradation resulting from PHD-bromodomain misfolding, consistent with what we observed in the bacterial overexpression experiment (Fig. 1e). The TRIM28-CRISPRi iPSCs displayed near-complete depletion of *TRIM28* mRNA and TRIM28 protein (Fig. S2h & Fig 2c).

**Fig. 2.**
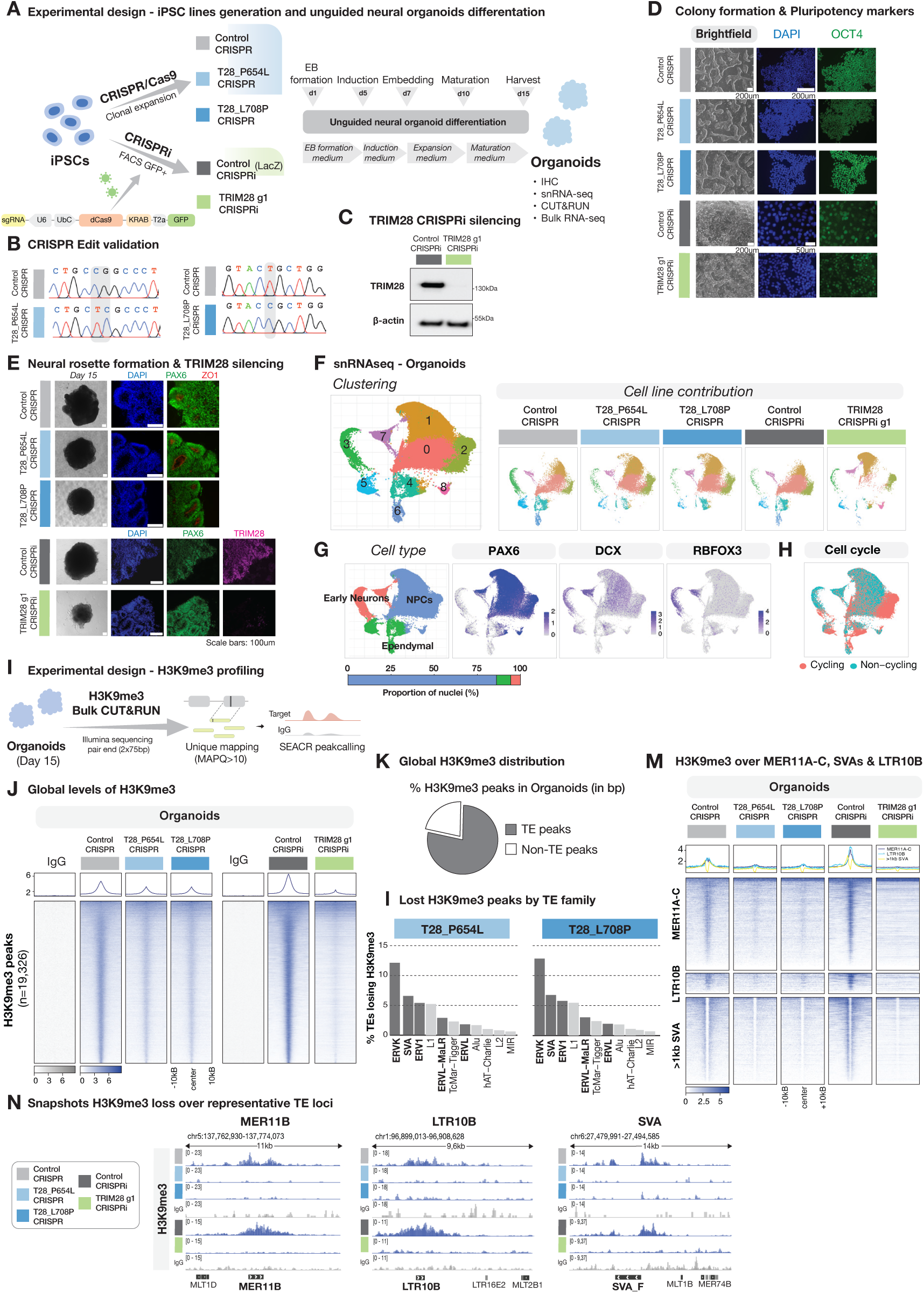
TRIM28 mutant organoids show loss of H3K9me3 loss over transposable elements. (**A**) Schematic of experimental design for generation of TRIM28 mutant and CRISPRi hiPSC lines, unguided neural organoid differentiation protocol and downstream analyses. (**B**) Chromatograms showing Sanger sequencing validation of the TRIM28 edits in the hiPSC lines. (**C**) Western blot (WB) of TRIM28 g1 CRISPRi hiPSCs protein extracts, TRIM28 (top) and B-Actin as loading control (bottom). (**D**) Column 1: Brightfield images showing colony formation and morphology of Control CRISPRi, P654L and L708P iPSCs. Column 2-4: Immunocytochemistry stainings against DAPI (blue) and OCT4 (green). (**E**) Column 1: Brightfield images of day 15 unguided neural organoids. Column 2-3: Immunohistochemistry against DAPI (Blue), PAX6 (green) and ZO1 (red) on day 15 unguided neural organoids. Column 4: Immunohistochemistry for DAPI (blue) and TRIM28 (magenta) on TRIM28g g1 CRISPRi and respective control day 15 unguided neural organoids. (**F**) Left: uniform manifold approximation and projection (UMAP) showing clusters found in day 15 unguided neural organoids. Right: UMAPs showing clustered nuclei per condition. (**G**) Left, top: UMAP displaying the different cell types found in day 15 neural organoids. Left, bottom: Bar plot showing the percentage cell type composition of the organoids at day 15 of differentiation. Right: UMAPs showing *PAX6*, *DCX* and *RBFOX3* expression as representative markers used to define specific cell types. (**H**) UMAP colored by cycling state. (**I**) Schematic of experimental design of H3K9me3 CUT&RUN profiling and bioinformatic approach. (**J**) Heatmaps illustrating genome-wide RPKM normalized CUT&RUN signal of non-targeting control IgG and H3K9me3 in Control CRISPR, T28_P654L CRISPR and T28_L708P CRISPR (n=2 per condition) (left) and Control CRISPRi and TRIM28 g1 CRISPRi (n=2 per condition) (right) day 15 unguided neural organoids. (**K**) Pie chart representing the percentage in base pairs of H3K9me3 peaks identified over TEs and non-TEs in day 15 unguided neural organoids. (**L**) Bar plots representing the percentage of TEs that lose H3K9me3 in the T28_P654L CRISPR (left) and T28_L708P CRISPR (right) mutant organoids sorted by family, highlighted in dark gray the elements belonging to the LTR and Retroposon families. (**M**) Heatmaps illustrating RPKM normalized CUT&RUN signal over TEs from the MER11A-C, SVAs and LTR10B subfamilies of non-targeting control IgG and H3K9me3 in Control CRISPR, T28_P654L CRISPR and T28_L708P CRISPR (n=2 per condition) (left) and Control CRISPRi and TRIM28 g1 CRISPRi (n=2 per condition) (right) day 15 unguided neural organoids. (**N**) Genome Browser tracks showing RPKM normalized H3K9me3 CUT&RUN data over representative MER11B, LTR10B and SVA elements.

To analyse the effects of the TRIM28 mutations on human brain development, we differentiated the edited hiPSCs into unguided neural organoids (Fig 2a). This model system allows us to study early human brain development in a 3D setting. We cultured organoids differentiated from one iPSC line from each condition for 15 days (Control CRISPR, P654L, L708P, Control CRISPRi and TRIM28-CRISPRi) (Fig. 2a). At this time point, the organoids represent an early stage of human brain development when neural progenitor cells (NPCs) are proliferating. All the iPSC lines could be successfully differentiated into neural organoids that formed neural rosettes, as monitored by ZO1/PAX6 immunostaining (Fig. 2e). We also confirmed TRIM28 silencing in the TRIM28-CRISPRi organoids after differentiation through immunostaining (Fig. 2e). To further investigate the cell-type composition, we analysed the organoids using single-nucleus RNA sequencing (snRNA-seq). High-quality data were generated from a total of 77,543 nuclei, including 15,878 from P654L organoids (15 organoids from 3 independent batches), 15,890 from L708P organoids (15 organoids from 3 independent batches) and 17,501 from TRIM28-CRISPRi organoids (16 organoids from 2 independent batches) as well as 13,293 and 14,981 nuclei from the two control groups (Control CRISPR: 11 organoids from 2 independent batches; Control CRISPRi: 16 organoids from 2 independent batches). We performed an unbiased clustering analysis to identify and quantify the different cell types present in the organoids. Nine separate clusters were identified, including neural cells at different stages of maturation (Fig 2f). These included proliferating and non-proliferating neural progenitor cells (NPCs) and newborn neurons, as well as cells with a transcriptional profile indicative of ependymal cells (Fig 2g&h, Fig. S2i). The contribution of the different cell lines to the clusters and cell types did not appear to differ substantially (Fig 2f, Fig. S2j), confirming that hiPSCs carrying *TRIM28* variants can successfully differentiate into neural cell types, such as NPCs and immature neurons.

### TRIM28 mutant neural organoids display a loss of H3K9me3 at transposable elements

To investigate the effect of the p.P654L and p.L708P missense variants in TRIM28-mediated H3K9me3 deposition, we performed CUT&RUN epigenomic profiling of H3K9me3 in iPSCs and day 15 unguided neural organoids (Control CRISPR, P654L, L708P, Control CRISPRi and TRIM28-CRISPRi, n=2 for each condition and each cell line). Because of the high degree of sequence similarity among TEs, short-read sequencing data results in a large proportion of reads that map ambiguously. To avoid signal amplification due to multimapping artifacts over these elements, we examined individual TE loci using a strict unique mapping approach (Fig. 2i). This bioinformatic approach allows us to analyze the epigenetic status of most TEs except for some of the evolutionarily youngest elements, (e.g. L1HS and SVAs) for which we rely on flanking regions. This approach enables us to distinguish reads unambiguously and trace epigenetic modifications to a specific TE locus in the human genome.

The CUT&RUN analysis revealed that H3K9me3 is abundant in both hiPSCs and neural organoids. Peak calling identified 11,383 H3K9me3-covered regions in hiPSCs and 19,362 H3K9me3 regions in neural organoids (Fig. S3a, Fig 2j). In both hiPSCs and neural organoids, most peaks could be confidently mapped to TEs, 70.2% in hiPSCs and 79.8% in neural organoids (Fig. S3b, Fig 2k). We found a genome-wide loss of the majority of H3K9me3 peaks in iPSCs and neural organoids carrying the P654L and L708P mutation, as well as in TRIM28-CRISPRi iPSCs and neural organoids (Fig. S3a, Fig 2j).

Since H3K9me3-associated heterochromatin was predominantly found over TEs (Fig. 2k), we further investigated which types of TEs lost this repressive mark in the *TRIM28* mutant hiPSCs and neural organoids. We found that several different types of TEs, such as ERVs, L1s, and SVAs, lost H3K9me3 both in iPSCs and neural organoids carrying the p,P654L and p,L708P mutation. In particular, certain classes of ERV elements and SVAs displayed a significant loss of H3K9me3, consistent with previous data on TRIM28-controlled TEs in human cells^31^ (Fig 2l). For instance, MER11A-C, LTR10B, and full-length SVAs showed a global decrease in H3K9me3 in both TRIM28 mutants and TRIM28 CRISPRi neural organoids (Fig. 2m&n). It has been shown in other models, including human NPCs, that TRIM28 is recruited by ZNF91 to SVA elements, leading to H3K9me3 deposition over these repetitive elements^11,45^. MER11A-C elements can be bound by several KRAB-ZFPs, like ZNF578 and ZNF808^22,23^. Interestingly, both transcription factors are able to recruit and bind TRIM28^24^ and are highly expressed in our unguided neural organoids. Together, these results demonstrate that these PHD-bromodomain mutations in TRIM28 result in loss-of-function effects, which are characterized by an absence of H3K9me3 deposition at TEs, particularly over ERVs and SVAs. This shows that particular TRIM28 missense variants can impair normal TRIM28 activity and have the potential to cause alterations in the epigenetic landscapes of pluripotent and differentiated cells.

### TRIM28 mutant organoids display transcriptional alterations in genes linked to neuronal differentiation and maturation

To investigate the impact of the p.P654L and p.L708P missense TRIM28 mutations on neural differentiation, we analyzed the snRNA-seq dataset from the neural organoids (Fig 2f). We focused on the NPC-like populations (Fig 2f), as they were the most abundant cell type in the neural organoids, and transcriptional changes in this population are likely to have functional consequences during subsequent differentiation into more mature cell types. To improve the resolution of the analysis of different NPC subpopulations, we subclustered the NPC snRNA-seq data (from Fig 2f; Clusters 0, 1, 2 & 8). This resulted in eight distinct clusters (Fig 3a). Interestingly, we identified clusters that were highly enriched for nuclei of the control condition (clusters 0 & 7), some for nuclei from organoids carrying either missense mutation (clusters 1 & 2) and others enriched for nuclei from the CRISPRi organoids (clusters 3, 6 & 8) (Fig 3b). These differences between conditions suggest significant transcriptomic changes, since they could not be explained by cell cycle state or major differences in cell identity (Fig 3c, d). We could also confirm in the snRNAseq data that *TRIM28* expression was silenced in the CRISPRi condition (Fig 3e). These results demonstrate that different NPC populations are largely transcriptionally distinct based on TRIM28 mutation and abundance, highlighting the significant impact of TRIM28 on the transcriptomic profile of NPCs in neural organoids.

**Fig. 3.**
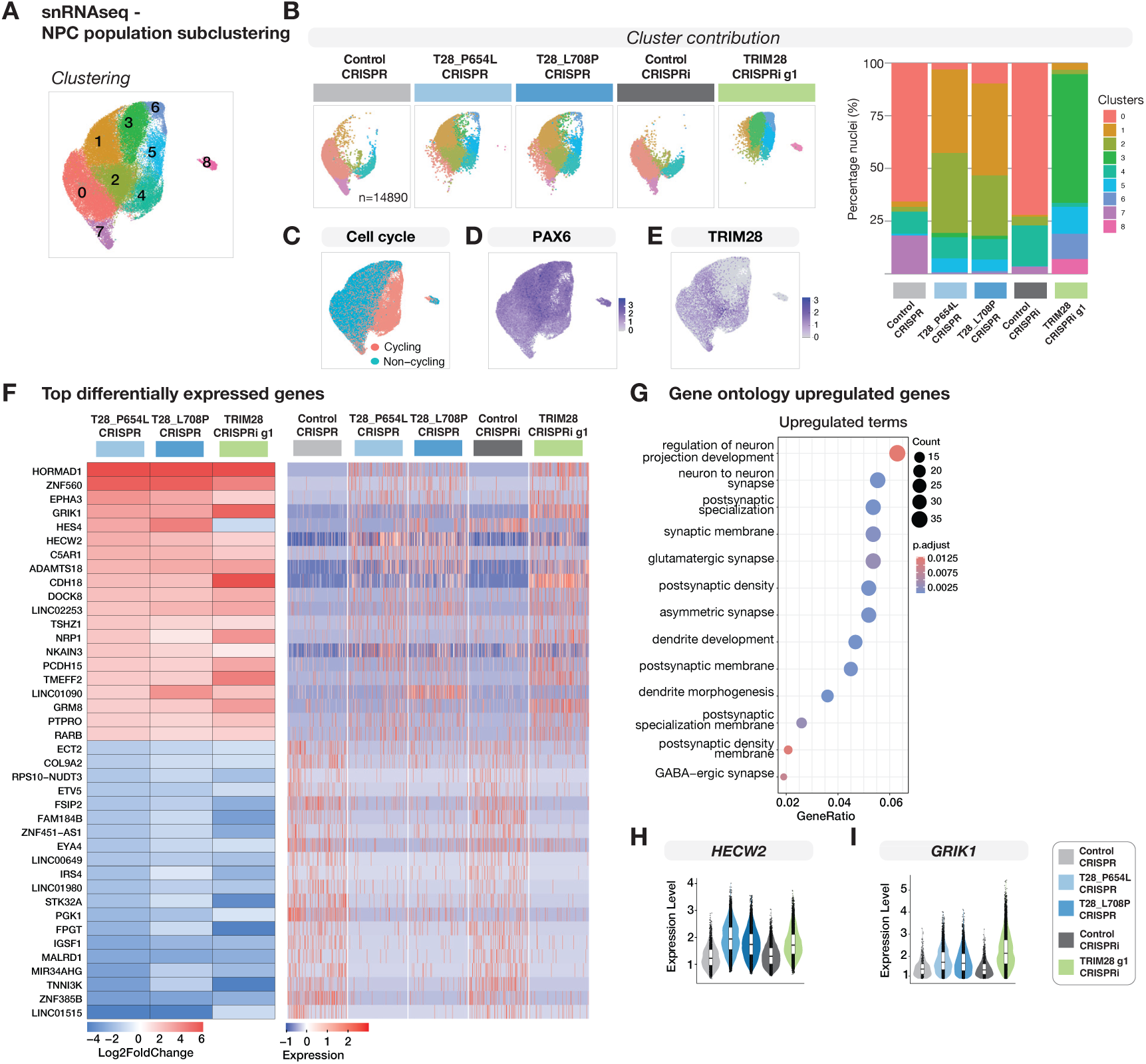
Transcriptional dysregulation in TRIM28 mutant neural progenitor cells. (**A**) UMAP showing subclustering of NPC population coloured by identified clusters. (**B**) Left: UMAPs showing NPCs clusters split by cell line. Right: Bar plot quantifying cluster contribution per cell line. (**C**) UMAP coloured by cycling state. (**D**) UMAP showing PAX6 expression. (**E**) UMAP showing TRIM28 expression. (**F**) Left: Heatmap showing the average Log_2_Fold change of top 20 up and downregulated genes in the P654L mutant in all conditions vs their respective controls. Right: Heatmap showing the same genes in the snRNA-seq data in all conditions. The full lists of differentially expressed genes in all conditions can be found as Supplementary Material. (**G**) Gene set enrichment analysis of upregulated genes in P654L NPCs compared to Control CRISPR. (**H**) Violin plots of *HECW2* expression in P654L, L708P and TRIM28-CRISPRi NPCs compared to their respective controls (Wilcoxon test). (**I**) Violin plots of *GRIK1* expression in P654L, L708P and TRIM28-CRISPRi NPCs compared to their respective controls (Wilcoxon test).

To explore the underlying transcriptional differences, we performed a differential expression analysis of the different TRIM28 perturbations against their respective controls. This analysis revealed reproducible transcriptional changes that were shared by the organoids obtained from the two mutant cell lines and the TRIM28-CRISPRi organoids. For example, we identified 578 genes that were significantly upregulated and 1225 genes that were downregulated in P654L NPCs (p-value < 0.05, log₂ fold change [LFC] > 0.5). Most of these genes were up– and downregulated in both the L708P condition and the TRIM28-CRISPRi organoids (Fig. S4a). Plotting the top 20 up– and downregulated genes across all conditions revealed that the differentially regulated P654L genes exhibited a similar pattern of transcriptional changes in L708P neural organoids and TRIM28-CRISPRi neural organoids (Fig. 3f). Similar results were obtained when the L708P mutation was used as the starting point for the analysis (Fig. S4b). The TRIM28-CRISPRi organoids shared most of the transcriptional changes observed in organoids with the two missense mutations but also displayed additional alterations likely resulting from TRIM28 functions independent of SETDB1 interaction (Fig. S4a).

Gene set enrichment analysis revealed that upregulated genes in the TRIM28 mutant NPCs were associated with neural specific terms, such as neuronal development, maturation, and synapse formation (Fig 3g), while the downregulated genes were related to more general cell homeostasis terms (Fig. S4c). For instance, *HECW2*, a ubiquitin protein ligase associated with epilepsy and intellectual disability^46^, was robustly up-regulated in organoids from the two mutant lines and the TRIM28-CRISPRi organoids (Fig 3h). Similarly, *GRIK1*, which encodes the glutamate ionotropic receptor kainate type subunit 1, was robustly upregulated in organoids from the two mutant lines and the TRIM28-CRISPRi organoids (Fig. 3i). Alterations in glutamate receptors have been associated with various neurodevelopmental and neuropsychiatric disorders^47^. In summary, these results demonstrate that neural organoids generated with the two mutant lines exhibit robust and consistent transcriptional changes. These changes are characterized by increased expression of genes associated with neuronal differentiation, which is largely reflected in the TRIM28-CRISPRi neural organoids. These results provide additional evidence that the two missense mutations in the PHD-bromodomain are loss-of-function alleles.

### Loss of TRIM28-mediated H3K9me3 deposition activates the regulatory potential of transposable elements

TEs carry regulatory sequences that can mediate *cis*-acting transcriptional effects on the surrounding genome. We therefore investigated whether the loss of TRIM28-mediated heterochromatin domains over TEs in neural organoids underlies the transcriptional changes observed in P654L and L708P mutant neural organoids (Fig 4a). Investigating the transcriptional changes of genes monitored by bulk RNA-seq in the mutant neural organoids and the TRIM28-CRISPRi organoids revealed profound *cis*-effects of TEs on nearby gene expression. The expression of genes located within 50 kb of a MER11A-C, LTR10B, or full-length SVA element was significantly increased in neural organoids generated from the two mutant iPSC lines, as well as in TRIM28-CRISPRi organoids (Fig 4b).

**Fig. 4.**
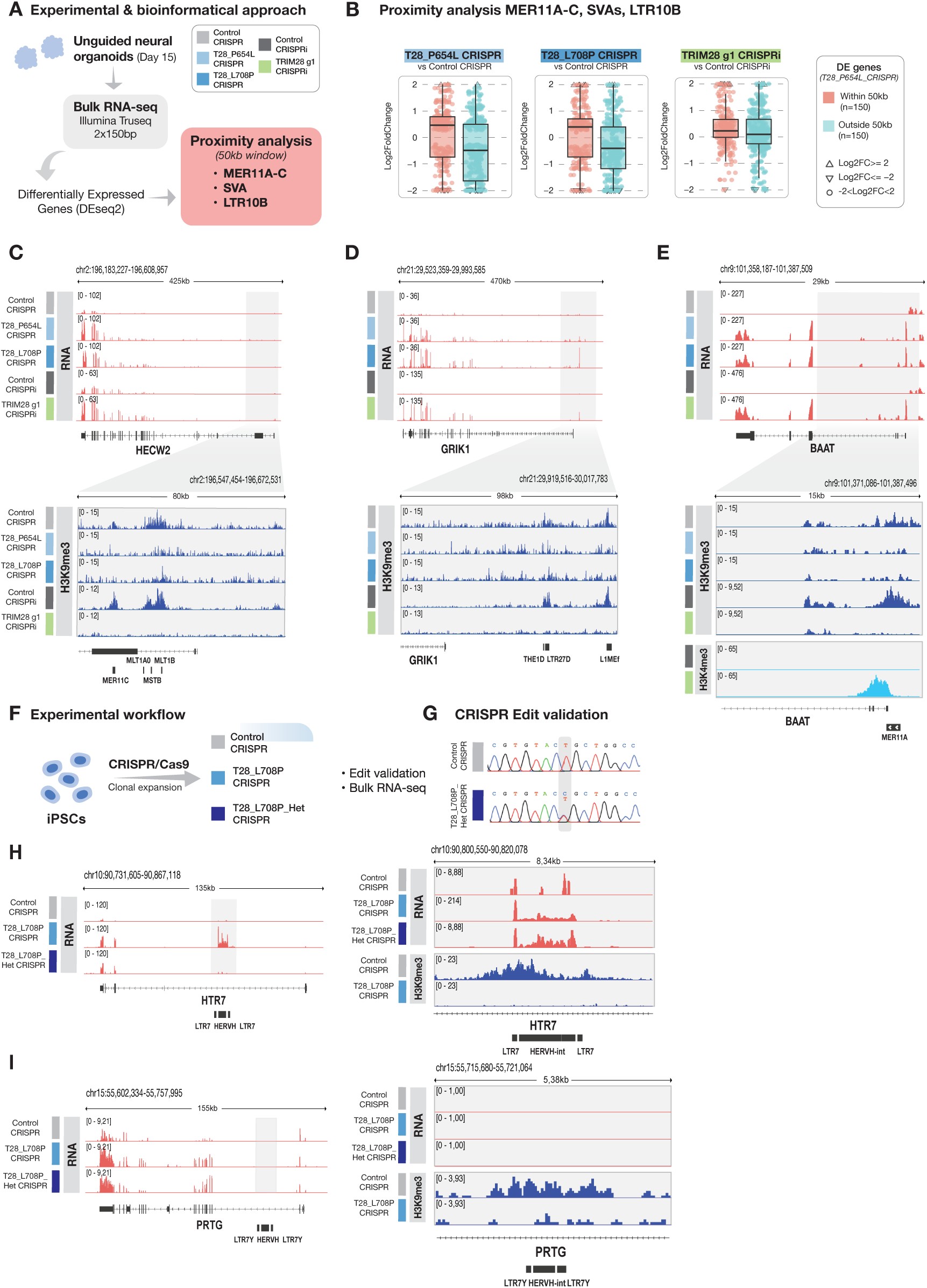
Cis-acting transcriptional effects in TRIM28 mutant organoids. (**A**) Schematic of experimental design of bulk RNA sequencing and bioinformatic approach for the proximity analysis. (**B**) Boxplots representing Log_2_FoldChange of P654L differentially expressed genes with a proximal MER11A-C, LTR10B or SVAs element (n=150; padj < 0.05; gene TSS within 50kb up/downstream of TEs), plotted next to differentially expressed genes without a proximal element (n=150; padj < 0.05; gene TSS outside 50kb window). Plotted Log_2_FoldChanges of the two gene lists for each perturbation – control pair. Statistical significance calculated with Wilcoxon test. (**C**) Above: Genome browser tracks showing bulk RNA sequencing signal over the *HECW2* locus. Below: Zoom-in of highlighted grey area showing RPKM normalized CUT&RUN enrichment profiles for H3K9me3. (**D**) Above: Genome browser tracks showing bulk RNA sequencing signal over the *GRIK1* locus. Below: Zoom-in of highlighted grey area showing RPKM normalized CUT&RUN enrichment profiles for H3K9me3. (**E**) Above: Genome browser tracks showing bulk RNA sequencing signal over the *BAAT* locus. Below: Zoom-in of highlighted grey area showing RPKM normalized CUT&RUN enrichment profiles for H3K9me3 and H3K4me3. (**F**) Schematic workflow for generation of L708P heterozygous clones. (**G**) Chromatograms showing Sanger sequencing validation of the TRIM28 edits in the hiPSC lines. (**H**) Left: Genome browser tracks showing bulk RNA sequencing signal over the *HTR7* locus. Right: Zoom-in of highlighted grey area showing bulk RNA sequencing signal and RPKM normalized CUT&RUN enrichment profiles for H3K9me3. (**I**) Left: Genome browser tracks showing bulk RNA sequencing signal over the *PRTG* locus. Right: Zoom-in of highlighted grey area showing bulk RNA sequencing signal and RPKM normalized CUT&RUN enrichment profiles for H3K9me3.

For example, we observed that *HECW2*, which is overexpressed in mutant organoids, has several LTR elements inserted within its introns located just downstream of its transcription start site (TSS) and that *GRIK1*, which is also overexpressed in mutant organoids, also has TE insertions located just upstream of its TSS. These TEs are covered by H3K9me3 in control organoids, but this repressive mark is absent over these elements in the two mutant neural organoids and in TRIM28-CRISPRi organoids (Fig 4c&d). These results suggest that TRIM28-mediated TE repression directly affects the expression levels of nearby genes, including those important for neural development. Additionally, we found genes located close to TEs that were activated in TRIM28-mutant organoids, and which are not typically expressed in neural tissues. For example, we identified an MER11A element that loses H3K9me3-associated heterochromatin in both TRIM28 mutant and TRIM28 CRISPRi neural organoids in the *BAAT* gene (Fig 4e). This gene encodes a hepatic enzyme that is exclusively expressed in the liver (GTEx V10, dbGaP accession phs000424.v10.p2)^39^. Taken together, these results suggest that loss of H3K9me3 over TEs directly affects the expression levels of nearby genes, including those important for neural development and those that should be silenced in neural tissues. Notably, we found little evidence of TE transcriptional activity itself in TRIM28 mutant organoids, suggesting that the main consequence of the loss of H3K9me3 over TEs is the activation of their gene regulatory potential.

### Dysregulation of TE gene networks in heterozygous L708P mutant iPSCs

Our CRISPR modelling experiments demonstrate that the p.P654L and p.L708P missense variants are loss-of-function alleles. However, in patients, these mutations are heterozygous. To investigate whether we can detect consequences of carrying these mutations in a heterozygous state, we generated iPSCs with a heterozygous p.L708P mutation (Fig 4f&g, Fig. S5a). Unfortunately, we were unable to generate p.P654L heterozygous clones. When we performed RNA-seq analysis on the heterozygous p.L708P iPSCs, we found that the consequences of this mutation were very modest overall in terms of the transcriptome (Fig. S5b). However, we did observe 11 commonly upregulated genes with the p.L708P homozygous line, albeit the effect was more modest in most cases. Interestingly, some of these commonly upregulated genes had a nearby TE that lost H3K9me3 in the p.L708P homozygous line. For example, in the *HTR7* gene, which encodes a serotonin receptor, we found an ERV that was epigenetically derepressed and transcriptionally activated in the homozygous p.L708P iPSCs and impacted the host gene. This effect was also present in heterozygous p.L708P iPSCs, albeit at a lower level (Fig. 4h). The *PRTG* gene, which encodes a member of the immunoglobulin superfamily that prevents precocious neuronal differentiation during embryonic neurogenesis^48^, was upregulated to a similar level in p.L708P homozygous and heterozygous iPSCs. This gene has an endogenous retrovirus (ERV) in its second intron that also loses H3K9me3 in the homozygous line. In this case, however, we could not detect transcriptional activation of the element in any of the genotypes (Fig. 4i). In summary, these results demonstrate that the effects of heterozygous missense mutations in the PHD-bromodomain of TRIM28 are modest but detectable. This lends further support to the hypothesis that *de novo* missense TRIM28 mutations is the underlying cause of the neurodevelopmental delay observed in these individuals.

## DISCUSSION

Several recent studies have demonstrated that transposable elements play an important role in controlling gene regulatory networks during brain development^10,11,40^. However, it has been unclear whether these observations are relevant to pathological consequences in humans. In this study, we demonstrate that *de novo* missense mutations in the master TE regulator TRIM28 result in neurodevelopmental delay characterized by dysregulation of TE-based gene regulatory networks. Our findings highlight the critical importance of the appropriate control of TEs during human brain development.

We identified two individuals with neurodevelopmental delay carrying *de novo* missense variants in *TRIM28*. These variants were found in the PHD-bromodomain, a protein domain that plays a critical role in the repressive actions of TRIM28. There is strong evidence to suggest that these missense mutations are the underlying cause of the neurodevelopmental syndrome reported in this study. *TRIM28* is highly expressed during human brain development, and data from mouse models indicate that it plays an important role in this process. Homozygous deletion of *TRIM28* during embryonic brain development is lethal^49^, while a heterozygous deletion results in animals with behavioral changes characterized by hyperactivity^34^. In addition, deletion of *TRIM28* in post-mitotic forebrain neurons in mice results in complex behavioural changes^35^ and heterozygous germ line deletion of *TRIM28* has been described to result in abnormal exploratory behaviour^36^. Furthermore, human population genetic data demonstrate that the PHD-bromodomain, in particular, is highly intolerant to mutations. Additionally, our induced pluripotent stem cell (iPSC)-based modeling using neural organoids demonstrates that the two PHD-bromodomain variants are loss-of-function alleles. This is evidenced by the loss of H3K9me3 and the transcriptional changes in organoids, including the disruption of TE-controlled transcriptional regulatory networks. These changes are largely mirrored by TRIM28-CRISPRi neural organoids, which represent a *bona fide* loss-of-function model and hiPSCs carrying heterozygous *TRIM28* variants, representing the patient situation, exhibit comparable transcriptional alterations albeit at a more modest level. In addition, some missense variants in this region (p.Leu708Pro, p.Arg767His, p.Pro754Leu) as well as other missense variants in *TRIM28* (p.Arg278Cys and p.Ser364Phe) have been reported *de novo* in individuals affected by developmental delay or autism included in the SPARC and ASC cohorts (Table S1)^50–52^. Taken together, these data strongly support the hypothesis that the *de novo* missense *TRIM28* variants in the PHD-bromodomain are the underlying cause in two individuals with a neurodevelopmental delay identified in this study and warrants the search for additional such cases.

It is well established that heterozygous mutations in *TRIM28* predispose individuals to Wilms’ tumor, a rare pediatric kidney cancer^53,49^, although no missense variants in the PHD-bromodomain have been reported in this context^54,55^. Approximately 1% of isolated and 8% of familial Wilms’ tumor patients harbor a germline variant in *TRIM28,* predominantly protein truncating or splice site disrupting mutations, suggesting complete loss of protein function^51^. Tumor development in these individuals follows a classic two-hit mechanism, with somatic inactivation of the remaining wildtype *TRIM28* allele, most frequently through copy-number neutral loss of heterozygosity in the developing kidney^56^. However, the published germline TRIM28 cohorts were ascertained through oncology settings and do not systematically report neurodevelopmental features in variant carriers, and there is no information available on the correlation between specific variant class and any neurodevelopmental phenotype. The low frequency of germline *TRIM28* variants in Wilms tumor (1-8% of cases) further limits the power of existing Wilms tumor cohorts to detect such an association. Published reports of neurodevelopmental abnormalities among Wilms’ tumor survivors focus on the effects of chemotherapy and radiation and not on germline variation^57^, and neurodevelopmental manifestations in variant carriers may therefore not have been recognized, or may have been attributed to other causes. Nevertheless, there is also a possibility that there is a differential effect between TRIM28 haploinsufficiency caused by nonsense, frameshift and splice site variants (associated with Wilm’s tumor and mostly absent in the ASD/DD databases) and the effect of these missense variants.

Our data demonstrate that TRIM28 plays a pivotal role in human brain development by regulating the activity of TEs within gene regulatory networks. Most notably, we found that the loss of H3K9me3 at TEs is associated with increased transcriptional activity of nearby genes. In several cases, TRIM28 represses TE regulatory activity, thereby preventing the expression of genes that should not be expressed in the neural lineage. Thus, in these cases, TRIM28 protects the genome from the impact of these TEs. We also found evidence of instances in which TRIM28 repression of TEs appeared to be co-opted to modify gene regulatory activity. For example, we identified TEs located near the TSS of *HECW2* and *GRIK1* that seem to decrease the expression of these genes. These observations suggest that TRIM28-controlled TEs are incorporated into gene regulatory networks to optimize the expression levels of protein-coding genes. Our results support a dual role for TEs in the genome. On the one hand, they threaten gene regulatory networks by incorporating new regulatory units. TRIM28 represses this activity and preserves genome integrity. Conversely, this activity can be harnessed to create new gene regulatory elements. These findings contribute to the growing awareness that TEs play important roles in gene regulatory networks, and that alterations to this activity can contribute to pathological states^16,18^. However, it is possible that TRIM28 functions unrelated to transposable element regulation may also contribute to the observed phenotypes. TRIM28 has been shown to play an important role in the control of imprinted genes as well as in DNA damage regulation^58,59^. Both of these features are also highly relevant for pathological processes in the developing brain.

In summary, our findings suggest a critical role of TE regulation in human brain development, as supported by the identification of *de novo TRIM28* loss-of-function missense variants in individuals with neurodevelopmental delay. These results provide strong support for continued investigations into the role of TEs in disorders with a neurodevelopmental component, such as psychiatric disorders. TRIM28-based repression of TEs depends on KZNFs, which are a large family of DNA-binding transcription factors recruited to TEs via sequence-based motifs. Genetic changes in KZNFs and their TE-targets are currently largely overlooked in genetic screens due to the repetitive nature of these sequences. However, future genome-based analyses using long-read sequencing approaches should resolve these issues and could reveal genetic variants in both KZNFs and TEs that contribute to the aetiology of these disorders.

## MATERIALS AND METHODS

### Patients and Exome Sequencing

Two unrelated patients ascertained via GeneMatcher^60^ were included in this study. Written informed consent for genetic testing and research use of clinical and molecular data was obtained from all participants or their legal guardians, in accordance with local ethics committee approvals (Ethics Committee of Hospital Clinic of Barcelona 2011/6625) and the Declaration of Helsinki.

Patient 1 was referred to the Hospital Clinic of Barcelona for genetic evaluation. Prior genetic testing included Fragile X testing and array comparative hybridization (aCGH), which did not identify any pathogenic alterations. Following these negative results, the patient was enrolled in a clinical research cohort and underwent exome sequencing. Genomic DNA from the patient and both parents was sent to the CRG (Barcelona, Spain) for trio exome sequencing as previously described^61^. Briefly, exonic regions were captured with the Agilent V3 (Agilent Technologies, Santa Clara, CA) exome capture kit and TruSeq indexing, and sequenced on a HiSeq™ 2000 Sequencing System (Illumina, San Diego, California, USA), generating 75 bp paired-end fragments. Reads were aligned to human genome build GRCh37/hg19 and analyzed using the eDiVa pipeline. The variant’s presence and its *de novo* status were validated by Sanger sequencing.

Patient 2 was evaluated at a genetics clinic in Canada at the age of 10, presenting with global developmental delay and significant behavioral problems including autistic traits, violent behaviour, severe onychophagia and hair chewing. Initial testing involved genetic testing for Fragile X, ATRX and Coffin-Lowry syndromes, and microarray, and metabolic screen of plasma and urine organic acids, acylcarnitine profile and uring glycosaminoglycans, which were all normal. The patient was then referred to GeneDx (Gaithersburg, MD, USA) for clinical trio exome sequencing. Using genomic DNA from the proband and parents, the exonic regions and flanking splice junctions of the genome were captured using the Clinical Research Exome kit (Agilent Technologies, Santa Clara, CA) or the IDT xGen Exome Research Panel v1.0. Massively parallel (NextGen) sequencing was done on an Illumina system with 100bp or greater paired-end reads. Reads were aligned to human genome build GRCh37/UCSC hg19 and analyzed for sequence variants using a custom-developed analysis tool. Additional sequencing technology and variant interpretation protocol has been previously described^62^. The general assertion criteria for variant classification are publicly available on the GeneDx ClinVar submission page (http://www.ncbi.nlm.nih.gov/clinvar/submitters/26957/)”

### Protein expression and solubility

Double-stranded gBlocks Gene Fragments (IDT) encoding the PHD-bromodomain (residues 624-812) wild-type sequence and p.P654L and p.L708P variants were cloned into expression vector pET-15b using NdeI and BamHI restriction sites. These constructs allowed production of N-terminally thrombin-cleavable His6-tagged protein products. Transformed E. coli BL21(DE3) cells (NEB) were grown at 37 °C in LB media containing 100 mg/L ampicillin. At an OD600 of ∼0.6, the uninduced sample (U) was collected and frozen and expression was induced with 0.2 mM IPTG for 3h at 37°C on a shaker, after incubation the induced sample (I) was collected. All subsequent steps were done at 4 °C unless otherwise stated. The culture was pelleted and resuspended in a lysis buffer containing 50 mM Tris pH 8.0, 300 mM NaCl, 10 mM imidazole, 1 mM DTT and 1× Roche complete EDTA-free protease inhibitors. Resuspended pellets were sonicated on in ice for 10 minutes (10s with 20s recovery cycles). A sample of this lysate was directly collected as the insoluble fraction (Ins), the rest was centrifuged for 20 minutes at maximum speed. The supernatant was then collected as the soluble sample (Sol). All samples were prepared for gel electrophoresis by adding 4x Bolt^TM^ LDS sample buffer (Invitrogen), 10x Bolt^TM^ sample reducing agent (Invitrogen) and 0,5ul benzonase nuclease (Sigma) and were boiled at 95°C for 5 minutes. Samples were loaded in a NuPAGE^TM^ 4-12% Bis-Tris mini protein gel (Invitrogen) and run for ∼30 min at 200V. The gel was stained with EZBlue^TM^ gel staining reagent (Sigma) for 2 hours, rinsed with deionized water and imaged.

### iPSC lines and maintenance

CTR14 human induced pluripotent stem (iPS) control cells (from Karolinska Institute iPS Core Facility) were reprogrammed from human fibroblasts from a young female donor using mRNA mediated reprogramming as described in Uhlin et al, 2017 (PMID 28395796) and under Ethics Review Board, Stockholm, March 28, 2012 (Registration number: 2012/208–31/3).

The iPSC cells were cultured on Laminin-521 coated (0,7 mg/cm2; Biolamina) Nunc plates in iPSC media (StemMacs iPS-Brew XF with 0,5% penicillin/streptomycin (GIBCO)). The iPSCs where passaged when 70-90% confluent as single cells with Accutase (GIBCO) and seeded in media supplemented with 10uM Y27632 Rock Inhibitor (Miltenyi Biotech). The media was changed daily. Cells were periodically tested for mycoplasma.

### CRISPR approaches

#### CRISPRi

To silence transcription of TRIM28 we used a CRISPR interference (CRISPRi) approach described elsewhere^11^. Lentiviral deadCas9-KRAB-T2A-GFP vectors including two different gRNA sequences targeting TRIM28 were used to silence TRIM28 expression. A control vector expressing a gRNA absent in the human genome (LacZ) was used as control. The single gRNA sequences used are listed in Table 2. Lentiviral vectors were produced as described below and yielded titers of 108-109 TU/ml determined by qPCR. An MOI of 15 was used in all conditions. On day 10 after transfection, GFP+ cells were FACS sorted (FACSAria, BD sciences) at 10°C. Sorted cells were pelleted at 1000 g at 10°C for 15 minutes, snap frozen on fry ice and stored at –80°C until RNA isolation. Knock-down efficiency was validated through qRT-PCR, RNA sequencing and western blot.

**Table 2.**
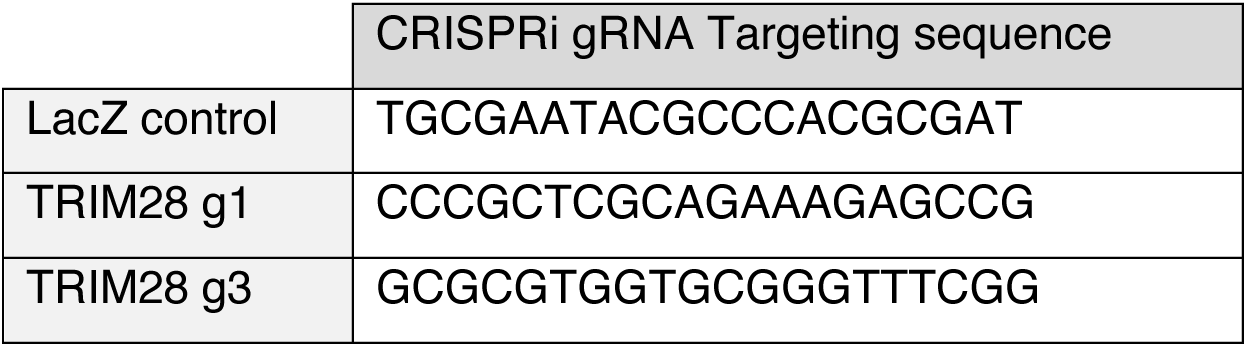
Target sequences of CRISPRi gRNA.

#### CRISPR editing & Quality controls

To introduce the patient mutation in the CTR14 iPSC line we used CRISPR editing. This was done by the Gene & Cell Therapy core at Lund University. The HiFi Cas9v3 (1081061), CRISPR-Cas9 tracrRNA (1072532) and edit specific crRNAs and ssODNs were purchased from IDT. The crRNAs and ssODNs sequences are listed in Table 3. The RNP and ssODN were delivered by nucleofection using the P3 Primary Cell 4D-Nucleofector X Kit (Lonza). Single cell dispension was achieved through limited dilution.

**Table 3.**
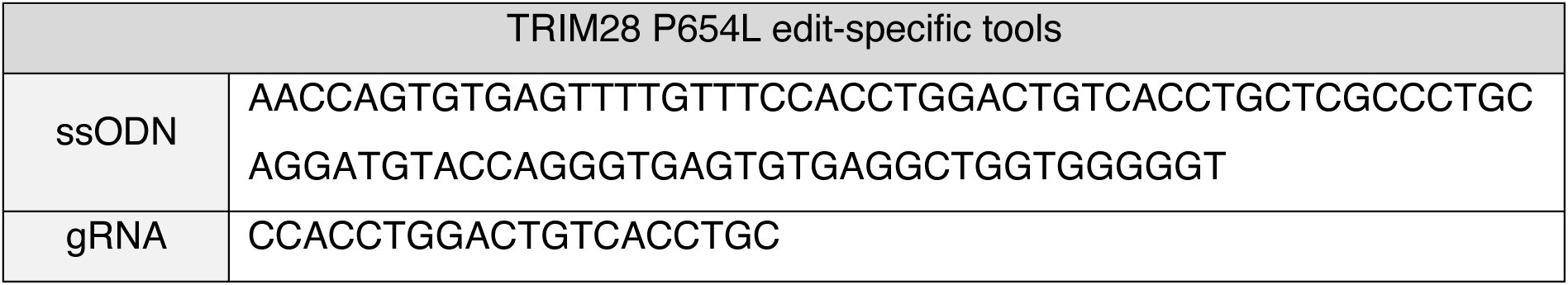
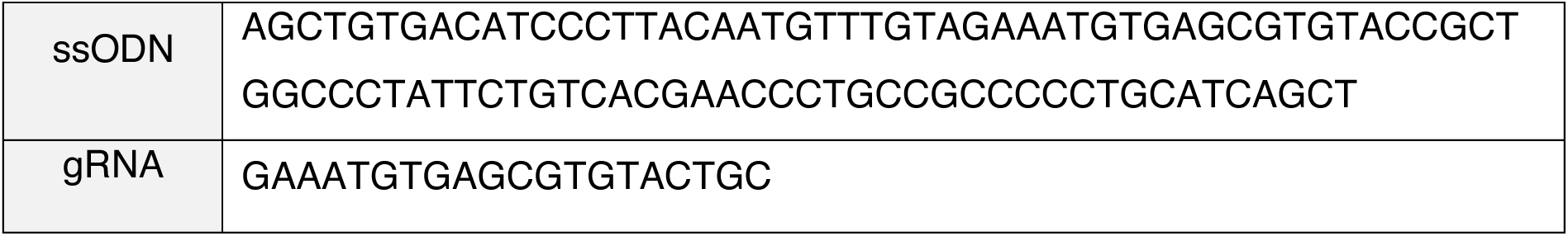
crRNAs and ssODNs sequences used in CRISPR editing.

The selected clones used in our experiments passed all the quality controls. These included: colony formation capacity; mycoplasma testing (Lonza, MycoAlertPLUS Mycoplasma Detection Kit), expression of pluripotency markers (see immunocytochemistry and antibodies sections for more detail), edit verification through Sanger sequencing (Eurofins Genomics, Mix2Seq Kit), cell line authentication based in STR analysis from genomic DNA (Eurofins Genomics, Cell Line Authentication Applied Biosystems AmpFLSTR Identifiler Plus PCR Amplification Kit system), karyotyping (Ambar Anàlisis Mèdiques) and validation of absence of edit in the top 5 off-target hits based on CFD score through Sanger sequencing (Eurofins Genomics, Mix2Seq Kit).

Quantitative genotyping PCR (qgPCR) was performed to rule out unintended CRISPR-induced on-target effects (onTEs) at the edited site. We followed the qgPCR protocol for detection of onTEs published by Weisheit et al., 2021. In brief, we extracted genomic DNA form iPSC pellets using the DNeasy Blood & Tissue kit (QIAGEN). Primers surrounding the edited regions and FAM-labeled target-specific probe were designed using the IDTPrimerQuest and purchased from IDT (Table 4). The TERT Taqman Copy Number Reference assay (Thermo Fisher Scientific) labeled with VIC was used as a control assay. The qgPCR reaction mix was prepared using the Prime Time Gene Expression Master Mix (IDT) and three technical replicates were included per sample. The reaction was run on a LightCycler 480 instrument (Roche). Analysis of Ct values and total allele number calculation were done according to the protocol guidelines.

**Table 4.**
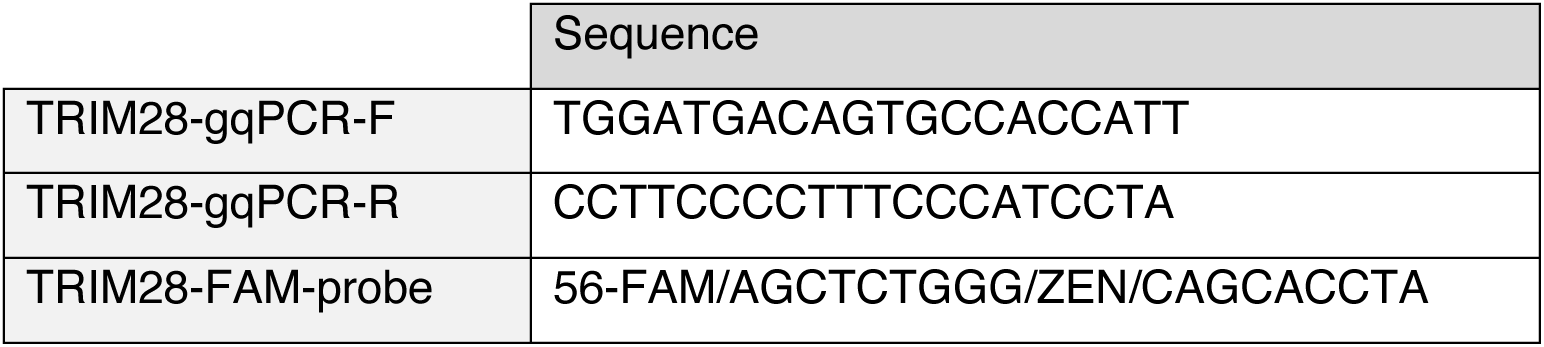
Sequences for amplification and detection of onTE in *TRIM28*.

#### Lentiviral production

Lentiviral vectors were produced according to a previously published protocol^63^. Briefly, two T175 Nunc flasks of 70-90% confluent HEK293T cells were transfected with the plasmid of interest and third-generation packaging and envelop plasmids (pMDL, psRev and pMD2G) using polyethyleneimine (PEI; Polysciences PN 23966) in DPBS (GIBCO). Two days after transfection, the supernatant was harvested, filtered, and centrifuged at 25000 g for 1,5 hours at 4°C. Lentiviruses were then resuspended in cold DPBS (GIBCO) and left at 4°C overnight. Resuspended lentiviruses were aliquoted and stored at –80°C.

#### Organoid differentiation

Unguided cerebral organoids were generated using the STEMdiff Cerebral Organoid Kit (STEMCELL Technologies) following manufacturer’s instructions. Briefly, 70-90% confluent iPSCs were detached using Accutase (GIBCO), resuspended in embryoid body (EB) formation media and counted. A total of 9,000 cells were seeded per well in U-bottom ultra-low attachment 96-well plates (Costar). The EBs were fed every two days until day 5 when they were transferred to ultra-low attachment 24-well plates (Sarstedt) containing neural induction media. Two days after, the EBs were embedded in Matrigel (Corning) and transferred to ultra-low attachment 6-well plates (Costar) with expansion media. On day 10, media was replaced with maturation media, then the organoids were fed every 3-4 days until harvested at day 15. We performed at least two independent batches of organoids for each experiment.

#### CUT&RUN

CUT&RUN experiments to profile histone modifications (H3K4me3, H3K9me3) were performed following a previously described protocol^64,65^. In brief, 500.000 cells per condition were harvested and washed 3 times (150mM NaCl, 0.5mM spermidine, 20uM HEPES pH 7.5, 1xRoche cOmplete protease inhibitors tablet) and incubated with ConA-coated magnetic beads (Bangs Laboratories) for 10 minutes at RT. Beads were pre-activated using binding buffer (20mM HEPES pH 7.9, 10mM KCl, 1mM MnCl2, 1mM CaCl2). The cells bound to the beads were the resuspended in antibody buffer (150mM NaCl, 20uM HEPES pH 7.5, 0.5mM spermidine, 1xRoche cOmplete protease inhibitors tablet, 0.05% w/v digitonin, 2mM EDTA) containing the appropriate primary antibody (see dilutions in antibodies section) and incubated at 4°C overnight with gentle shacking. The following day, beads were washed twice with digitonin buffer (150mM NaCl, 20uM HEPES pH 7.5, 0.5mM spermidine, 1xRoche cOmplete protease inhibitors tablet, 0.05% w/v digitonin). Beads were resuspended in a digitonin buffer containing pA-MNase (produced in-house) and incubated at 4°C for approximately 1 hour. After two more washes, beads were resuspended in 100 ul digitonin buffer and chilled to 0-2°C for 5 minutes. To activate genomic cleavage, 2mM CaCl2 were added to each sample and incubated at 0°C for 30 minutes. Finally, the reaction was quenched by adding 100 ul of 2x stop buffer (0.35M NaCl, 20mM EDTA, 4mM EGTA, 50 ng/ml RNase A, 50 ng/ml glycogen, 0.05% w/v digitonin) while vortexing gently. The samples were immediately incubated at 37°C for 30 minutes to release the genomic fragments from the insoluble nuclear chromatin. Beads were then removed using a magnetic stand and the DNA fragments from the supernatant were purified with a PCR clean-up spin column kit (Macherey-Nagel). For the organoid experiments, 6-8 organoids were collected on day 15 of differentiation, washed with PBS and incubated with Accutase (GIBCO) at 37°C on a shaker for 3 minutes, gently resuspended and incubated for 4 additional minutes. The cell suspension was then centrifuged at 400xg for 5 minutes and transferred to a Falcon™ Test Tube with Cell Strainer Snap Cap (Fisher Scientific) to remove Matrigel and achieve a single-cell suspension before continuing with the CUT&RUN protocol.

#### CUT&RUN sequencing and analysis

Sequencing libraries were prepared using the Hyperprep kit (KAPA) with unique dual indexed adaptors. Then they were pooled and sequenced using Illumina NovaSeqX plus with paired-end reads (2×75). Obtained reads were mapped to the human genome (GRCh38) using bowtie2 (v 2.4.5)^66^ (–local –very-sensitive-local –no-mixed –no-unal –no-discordant –phred33-I 10 –X 700), upon which they were converted BAMs and further sorted using SAMtools (v 1.16.1). Peaks were called with SEACR (v 1.3) using IgG samples as control for peak calling. Reads per kilobase per million mapped reads (RPKM)– normalized BigWig coverage tracks were made with bamCoverage (deepTools, v 2.5.4). Heatmaps matrices were created by computeMatrix (deepTools v2.5.4) using the called peaks as the input regions and RPKM normalized bigwigs as input signal. The matrices were visualized with plotHeatmap (deepTools, v2.5.4), adjusting the upstream and downstream visualization parameters (–a and – b) and splitting the signal in heatmaps (––kmeans 2).

#### Immunocytochemistry

The cells were grown on coated plastic coverslips or 24-well ibidi plates. When the desired confluence was reached, cells were rinsed three times with DPBS, fixed with 4% PFA for 15 minutes at room temperature and rinse three times more with DPBS. For blocking, they were incubated with 5% Normal Donkey Serum (NDS) in TKPBS (KPBS with 0,25% Triton X-100) for 1 hour at room temperature. Primary antibodies were then diluted in 5% NDS in TKPBS and incubated overnight at 4°C, no primary antibody was added to the negative controls. The following day cells were washed three times with TKPBS for 5 minutes and incubated for 2 hours at room temperature with the appropriate secondary antibodies (see below) diluted in 5% NDS in TKPBS. For nuclear counterstaining, cells were incubated with DAPI (Sigma D817; 1:1000) for 5 minutes. Two final 5 minutes washes with KPBS were done. Coverslips were rinsed with Milli-Q water, mounted with FluorSave (Millipore) on microscopes slides and imaged using a fluorescence microscope (Leica). Cells on ibidi plates were left with KPBS at 4°C until imaged using an Operetta CLS High Content Analysis imager (PerkinElmer) or a fluorescence microscope (Leica). Images were processed using ImageJ Fiji.

#### Immunohistochemistry

Organoids were collected at day 15, washed with DPBS and fixed with 4% PFA for 90 minutes at room temperature. They were then washed three times with KPBS and left in a 30% sucrose in PBS solution overnight at 4°C with gentle shaking. The fixed organoids were then transferred to a cryomold with OCT (HistoLab), frozen on dry ice and stored at –20°C until used. Before staining, the organoids were sectioned on a cryostat at a thickness of 20μm and placed on Superfrost plus slides. The slides were washed with KPBS for 5 minutes and blocked and permeabilized in 0,1% Triton X-100 and 5% NDS in KPBS for 1 hour at room temperature. The appropriate dilution of primary antibodies (see below) was prepared in blocking solution and incubated overnight at 4°C. The following day, the sections were washed three time with KPBS and incubated with the corresponding secondary antibodies diluted in blocking solution for 1 hour at room temperature. Finally, they were washed three times with KPBS, incubated with DAPI (Sigma D817; 1:1000) for 5 minutes and wash once more with KPBS. Slides were mounted with FluorSave (Millipore) and imaged using a fluorescence microscope (Leica). Images were processed using ImageJ Fiji.

#### Western blot

RIPA buffer (Sigma-Aldrich) supplemented with complete protease inhibitor cocktail (Roche) was used to lyse the cells on ice for 30 minutes. Lysed cells were then centrifuged at 12,000 rpm for 20 minutes at 4°C. Supernatants were collected, mixed with Bolt LDS loading buffer 4x (Novex) and Bolt reducing agent 10x (Novex) boiled for 5 min at 95°C and loaded in pre-made 4-12% Tris-glycine SDS-PAGE gels (Invitrogen). The extracts were run for 45 min at 200V, proteins were then transferred to a PVDF membrane using the Transblot-Turbo Transfer system (BioRad). Once transferred, the membrane was rinsed with Tris-buffer saline with 0,1% Tween (TBST) and blocked for 1-2 hours in TBST with 5% skimmed milk. Primary antibodies were diluted in TBST with milk at appropriate concentrations (see below) and incubated overnight at 4°C with gentle shaking. The day after the membrane was washed three time in TBST for 10 min and incubated for 1h at room temperature with HRP-conjugated anti-mouse secondary antibody (Cell Signaling) diluted in TBST with milk. After incubation, the membrane was washed twice with TBST and once with TBS before the protein was detected by chemiluminescence using ECL Select reagents (Cytiva) following manufacturer’s instructions on a Chemi-Doc system (BioRad).

#### Antibodies

Mouse anti-TRIM28 (Abcam, ab22553; 1:1000 dilution) and HRP-conjugated anti-b-actin (Sigma, A3854, 1:50000) were used for Western blots. Rabbit anti-H3K9me3 (Abcam, ab8898), rabbit anti-H3K4me3 (Active Motif, 39159) and goat anti-rabbit IgG (Abcam, ab97074) were used for CUT&RUN at a 1:100 dilution. Mouse anti-TRIM28 (as above; 1:100), mouse anti-OCT4 (Santa Cruz Biotechnology, sc-5279; 1:200), rabbit anti-Nanog (Abcam, ab21624; 1:200), rabbit anti-SOX2 (Merck AB5603; 1:500), rabbit anti-PAX6 (biolegend, Poly19013; 1:100), donkey anti-rabbit Alexa Fluor 488 (Thermo Fisher, AB_2535792; 1:500), donkey anti-mouse Alexa Fluor 488 (Thermo Fisher, AB_141607; 1:500) and donkey anti-mouse Alexa fluor 647 (Themo Fisher, AB_2762831; 1:500) were used for immunocytochemistry and/or immunohistochemistry.

#### RNA extraction and bulk RNA sequencing

RNA was extracted using the RNeasy mini kit and QIAshredder kits (QIAGEN) with on-column DNAse treatment for iPSCs and organoids, respectively. The isolated total RNA was used for RNA sequencing and/or RT-qPCR analysis (see below). The sequencing libraries were prepared using the Illumina Truseq Stranded mRNA library prep kit (poly-A selection) and sequenced on a NovoSeq X Plus with 150bp paired end reads.

*Gene quantification.* Reads were aligned to the human reference genome hg38 (GRCh38) using STAR aligner (v 2.6.0)^67^ and modified parameters (––outFilterMultimapNmax 100 and –-winAnchorMultimapNmax 200). BigWig files were created from the sorted BAM files with bamCoverage (v 2.5.4) applying RPKM normalization for visualization. Gene quantification was conducted with featureCounts (Subread v1.6.3) using GENCODE v40 as the reference annotation.

*Repetitive element quantification.* To perform accurate quantification of repetitive elements, only uniquely mapped reads were allowed when aligning with STAR (v 2.6.0c). Specifically, the default parameters were modified to only allow mapping to a single locus (––outFilterMultimapNmax 1) and the allowed mismatched ratio was decreased (––outFilterMismatchNoverLmax 0.03). Quantification was done as described above, except the curated annotation file provided within the TEtranscripts package (v 2.2.3)^68^ was used.

*Differential expression analysis.* Differential expression analysis was conducted with DESeq2(v 1.40.2)^69^ inputting the count matrices obtained from featureCounts. The raw reads were normalized with median-of-ratios normalization and fold changes were shrunk using DESeq2’s lfcShrink. When generating heatmaps, the mean of replicates for each condition was used along with “+0.5” pseudocount. The resulting data was visualized using pheatmap.

#### RT-qPCR

Isolated RNA was quantified using a Nanodrop 2000 (Thermo Fisher). 500 ng of RNA from each sample were reverse transcribed to cDNA using the Maxima First Strand cDNA Synthesis Kit (Thermo Fisher). The cDNA was analysed by RT-qPCR using SYBR Green I master (Roche) on a LightCycler 480 instrument (Roche) in triplicates. Representation of the data was done using the ΔΔCt method normalizing to GAPDH or YWHAZ as housekeeping genes. Primers sequences can be found below.

**Table 5.**
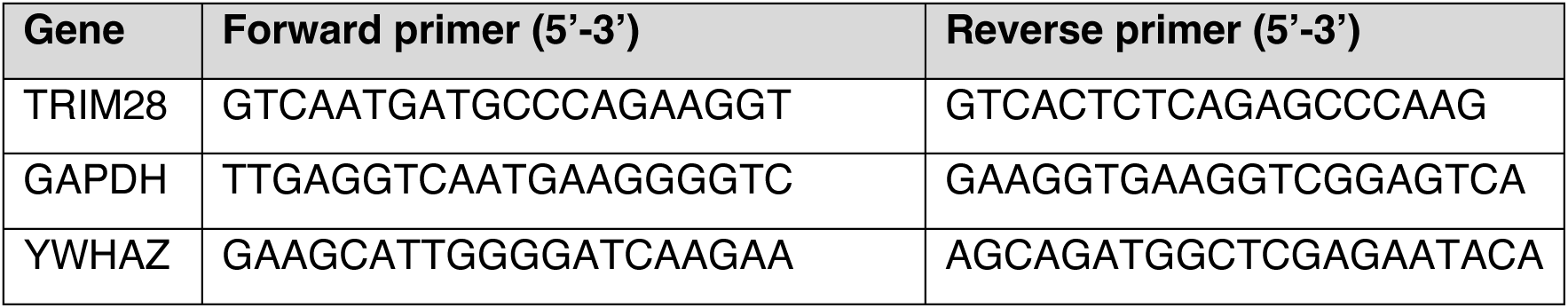
RT qPCR primers.

#### Nuclear isolation

Organoids were harvested for single nuclei RNA sequencing (snRNA-seq) at day 15. For collection, they were snap frozen on dry ice and stored at –80°C. Aiming to reduce possible biases caused by organoid inherent heterogeneity, each sample consisted of 4-5 organoids pooled together. We followed the nuclei isolation protocol using a sucrose gradient-based isolation previously described elsewhere^70^. In brief, the organoid samples were thawed and homogenized using a tissue douncer (Wheaton) in ice-cold lysis buffer (0.32M sucrose, 5mM CaCl2, 3mM MgAc, 0.1mM Na2EDTA, 10mM Tris-HCl pH 8.0, 1mM dithiothreitol). The homogenate was then carefully layered on a high sucrose solution (1.8M sucrose, 3mM MgAc, 10mM Tris-HCl pH 8.0, 1mM dithiothreitol) and centrifuged at 30,000xg for 2 hours and 15 minutes. The resulting pelleted nuclei were incubated for 10 min in 100ul of nuclear storage buffer (10mM Tris-HCl pH 7.2, 70mM KCl, 2mM MgCl2) and resuspended using a 70um cell strainer. Draq7-negative single nuclei were sorted at 4°C and low rate with a FACS Aria (BD Biosciences) using a 100um nozzle. Reanalysis showed purity of >95%.

#### Single-nuclei RNA sequencing and analysis

A total of 10,000 nuclei were sorted per sample and loaded onto the Chromium Next GEM Chip G Single Cell Kit together with the reverse transcription mastermix following manufacturer’s instructions for the Chromium Next GEM single cell 3’ kit (10X Genomics, PN-1000268) to generate an emulsion with single-cell gel beads (GEMs). Amplification of cDNA was performed according to 10x Genomics guidelines (13 cycles of amplification of the 3’ libraries) and sequencing libraries were generated with unique dual indices (TT set A) for each individual sample, pooled and sequenced on a Novaseq X Plus with a 100-cycle kit and 28-10-10-90 reads.

Cell Ranger count (v 6.0.0) was run with standard settings^71^, with an mRNA and pre-mRNA reference for single-cell samples for single-nucleus samples (based on 10x Genomics Cell Ranger 6.0.0 guidelines). Seurat (version 5.1.0)^72^ was then used to analyze sample clustering. Low quality nuclei from each sample were filtered out according two criteria; 1) percentage of mitochondrial content over 2% (perc_mitochondrial) and 2) number of features (n_Feature) lower than 600 and greater than 2 standard deviations. Seurat’s LogNormalize (NormalizeData) was used to normalize counts and clusters were defined with a resolution of 0.1 (FindClusters). All downstream visualization of the data was done with Standard Seurat functions.

## Acknowledgements

We would like to thank S. Henikoff for providing reagents. We thank J. G. Johansson, A. Hammarberg, S. da Rocha Baez, M. Persson-Vejgården, J. Nelander, B. Mattsson and U. Jarl for technical assistance. We acknowledge Clinical Genomics Lund, SciLifeLab, and Center for Translational Genomics (CTG) Lund University for providing expertise and service with sequencing and analysis, SCC Cell and Gene Therapy Core at Lund University (gene-editing service, www.stemcellcenter.lu.se/cellandgene) and KI iPS Core (https://ki.se/en/research/ips-core-facility) for assistance with iPSC culture and CRISPR-editing. We are grateful to all members of the Jakobsson laboratory. We also thank the families for their participation and the permissions to use the data of the patients.

## Funding

Swedish Research Council grant 022-00673 (JJ)

Swedish Research Council grant 2021-03494 (CD)

Swedish Brain Foundation grant FO2023-0232 (JJ)

Cancerfonden grant 222185 to (JJ)

Barncancerfonden grant PR2023-0099 (JJ)

Swedish Society for Medical Research grant S19-0100 (CD)

Chan Zuckerberg Initiative DAF grant 2023-331773 (CD)

Olle Engqvists stiftelse grant 218-0090 (JJ)

Lindhes Advokatbyrå grant LA2023-0077 (LCV)

Kungliga Fysiografiska Sällskapet i Lund (LCV)

Anna-Lisa Rosenberg stiftelse (LCV)

Thorsten och Elsa Segerfalks Stiftelse (LCV)

Swedish Government Initiative for Strategic Research Areas (MultiPark & StemTherapy)

Spanish MCIN /AEI/ 10.13039/501100011033 (SB, RR)

MCIU/AEI/ 10.13039/501100011033 and the ERDF “A way of making Europe” grants PID2019-17188RB-C21 and PID2022-141461OB-I00 (SB, RR)

AGAUR from the Autonomous Catalan Government grants 2021-SGR-01492 (IM) and 2021-SGR-1093 (RR)

CIBERER ACCI18-15-703 (RR)

## Author Contributions

Design and interpretation: All authors. Conceptualization: LCV, RR, JJ. Patient recruitment and evaluation: MIAM, IM, FBA, CP, SC. Experimental research: LCV, NP, CDH, FD, OK, CD. Bioinformatic analyses: NP, AMM, RR. Writing—original draft: LCV, NP, CDH, RR, JJ. Writing—review and editing: All authors.

## Competing Interests

AMM is an employee of and may own stock in GeneDx.

## Data Availability

The iPSC RNA and DNA sequencing data included in this study will be deposited at GEO upon publication. Previously published data used in this paper can be found at GEO Series GSE224747 (https://www.ncbi.nlm.nih.gov/geo/query/acc.cgi?acc=GSE224747).

## Supplementary Materials

**Supplementary Figure 1.**
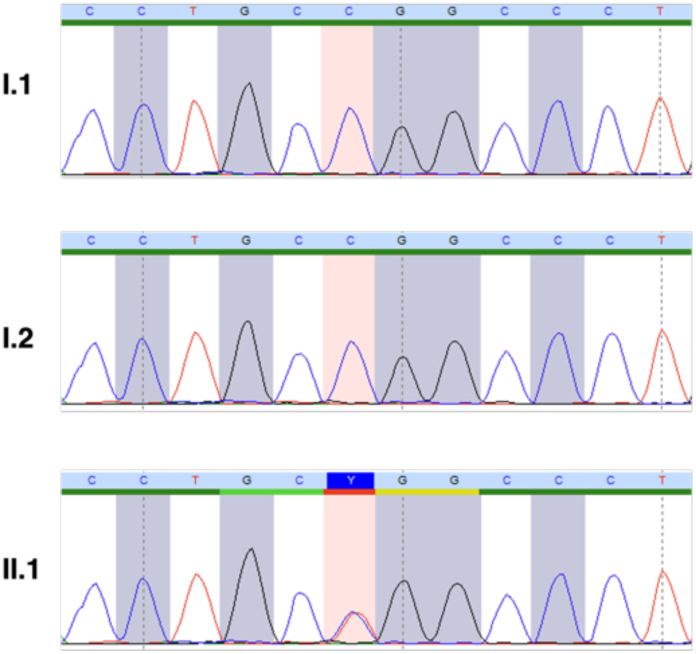
Chromatograms showing Sanger sequencing validation of the variant in patient 1 (II.2) and parents (I.1, I.2).

**Supplementary Figure 2.**
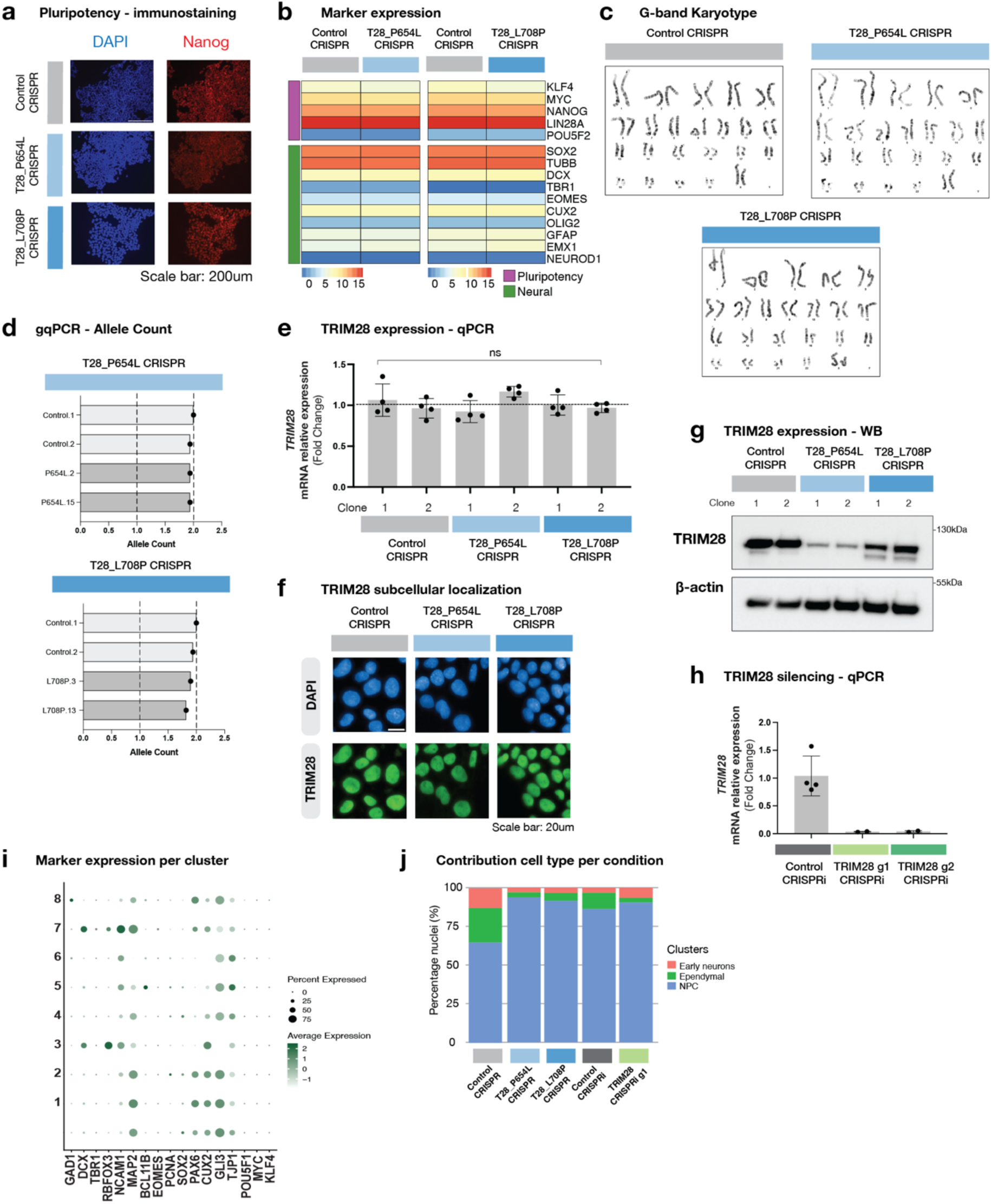
a) Immunocytochemistry staining against DAPI (blue) and Nanog (red) of Control CRISPRi, P654L and L708P iPSCs. b) Heatmap showing log2 normalized expression (RNA-seq) of pluripotency and differentiation markers in Control CRISPR vs. P654L and Control CRISPR vs. L708P hiPSCs. c) G-band karyotyping of Control CRISPRi P654L and L708P iPSCs clones. d) TRIM28 allele count genomic quantitative PCR of Control CRISPR, P654L and L708P iPSCs normalized to TERT. e) RT-qPCR analysis of TRIM28 expression in the TRIM28-Mut iPSCs (n=4 biological replicates) normalized to GAPDH. Bars represent mean ± SD. One-way ANOVA with Dunnett’s post hoc test for multiple comparisons. f) Immunocytochemistry stainings against DAPI (blue) and TRIM28 (green) of one representative Control CRISPRi P654L and L708P iPSCs. g) Western blot (WB) of TRIM28-Mut iPSCs protein extracts, TRIM28 (top) and B-Actin as loading control (bottom). Representative blot of 3 independent experiments. h) RT-qPCR analysis of TRIM28 silencing in the TRIM28 CRISPRi iPSCs (n=2 biological replicates) normalized to GAPDH. Bars represent mean ± SD. i) Dot plot displaying selected neuronal and NPC markers used to characterize the cell clusters (dot size shows the percentage of nuclei expressing the gene, color indicates average expression). j) Bar plot quantifying cell type contribution per cell line in unguided neural organoids.

**Supplementary Figure 3.**
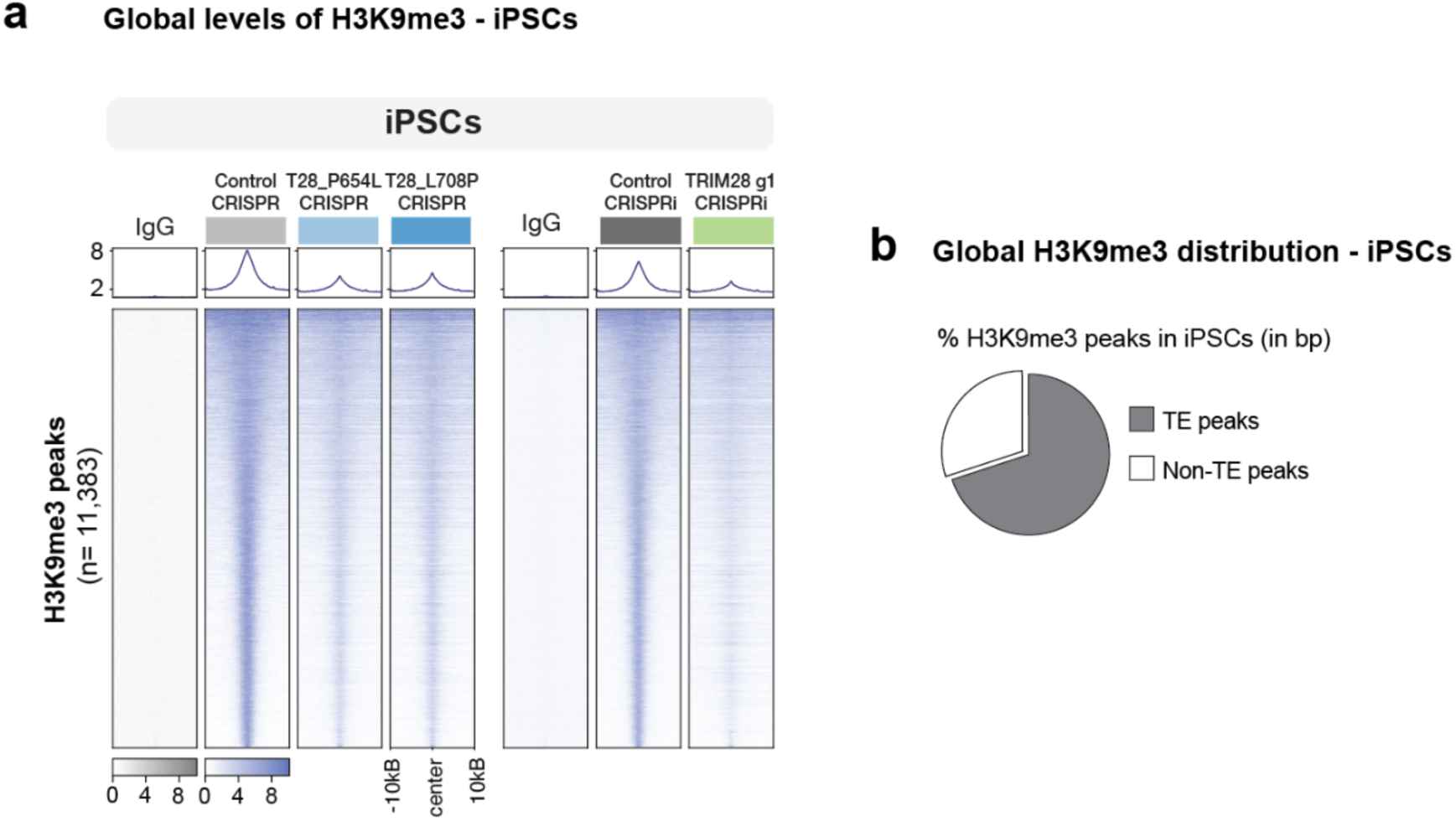
a) Heatmaps illustrating genome-wide RPKM normalized CUT&RUN signal of non-targeting control and H3K9me3 IgG in Control CRISPR, T28_P654L CRISPR and T28_L708P CRISPR iPSCs (n=2 per condition) (left) and Control CRISPRi and TRIM28 g1 CRISPRi iPSCs (n=2 per condition) (right). b) Pie chart representing the percentage in base pairs of H3K9me3 peaks identified over TEs and non-TEs in iPSCs.

**Supplementary Figure 4.**
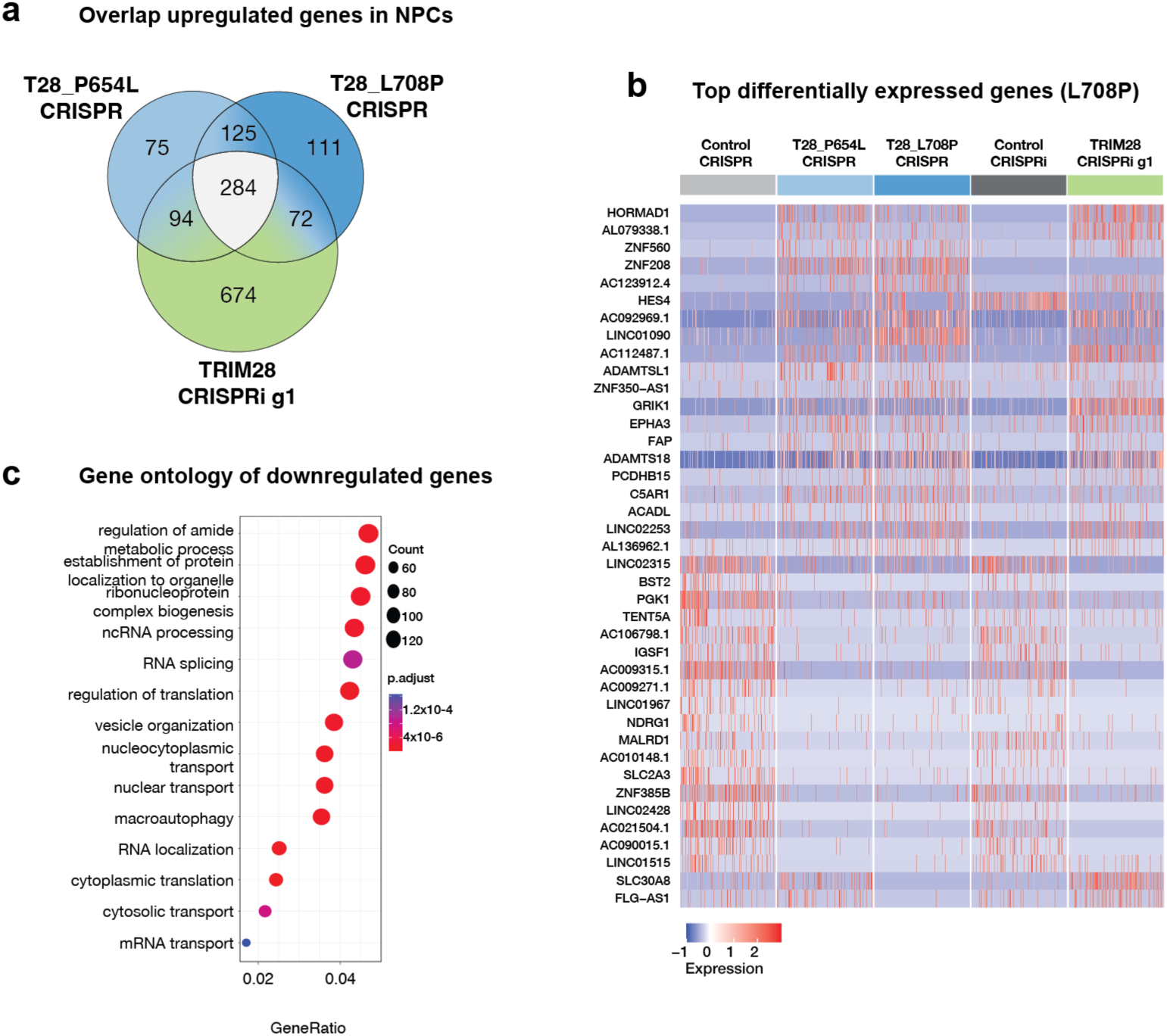
a) Venn diagram displaying common upregulated genes in the NPC population of T28_P654L CRISPR, T28_L708P CRISPR and TRIM28 g1 CRISPRi day 15 unguided neural organoids (against their respective controls). b) Heatmap showing the snRNA-seq data of top 20 up and downregulated genes on the L708P mutant in all conditions vs their respective controls. The full lists of differentially expressed genes in all conditions can be found as Supplementary Material. c) Gene set enrichment analysis of downregulated genes in P654L NPCs compared to Control CRISPR.

**Supplementary Figure 5.**
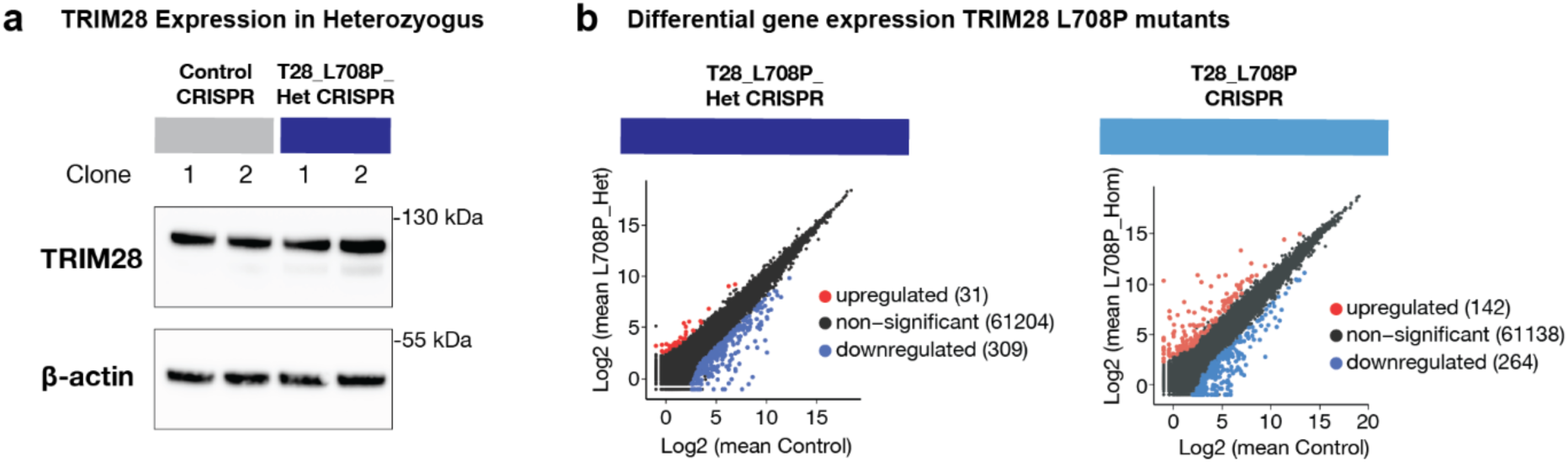
a) Western blot of T28_L708P_Het CRISPR and Control CRISPR protein extracts, TRIM28 (top) and B-Actin as loading control (bottom). Representative blot of 2 independent experiments. b) Mean plot showing differential gene expression in T28_L708P_Het CRISPR compared to Control CRISPR (n=4) and T28_L708P CRISPR compared to Control CRISPR (n=4) iPSCs. LFC >2, padj < 0.05 calculated with DESeq2.

**Supplementary Table 1:** *de novo TRIM28* missense variants in the literature associated with neurodevelopmental delay or autism.

| Variant | Sex | cDNA | Cohort | Alph-amissense | REVEL | CADD | Reference | Other |
| --- | --- | --- | --- | --- | --- | --- | --- | --- |
| R278C | Male | c.832C>T | ASCB14 | 0.7687 | 0.116 | 28.5 | 52 |  |
| S364F | Female | c.1091C>T | SPARK | 0.9996 | 0.408 | 25.4 | 51,52 |  |
| L708P | Male | c.2123T>C | GeneDX | 0.9993 | 0.483 | 25.7 | 50, 51, * |  |
| P754L | Male | c.2261C>T | SPARK | 0.7294 | 0.227 | 21.8 | 51,52 | Shared with unaffected sibling |
| R767H | Male | c.2300G>A | ASCB14 | 0.1715 | 0.037 | 26.9 | 51,52 |  |

\* it is possible that the previously published proband with this variant is the same individual reported in this manuscript

